# Favipiravir antiviral efficacy against SARS-CoV-2 in a hamster model

**DOI:** 10.1101/2020.07.07.191775

**Authors:** Jean-Sélim Driouich, Maxime Cochin, Guillaume Lingas, Grégory Moureau, Franck Touret, Paul Rémi Petit, Géraldine Piorkowski, Karine Barthélémy, Bruno Coutard, Jérémie Guedj, Xavier de Lamballerie, Caroline Solas, Antoine Nougairède

## Abstract

Despite no or limited pre-clinical evidence, repurposed drugs are massively evaluated in clinical trials to palliate the lack of antiviral molecules against SARS-CoV-2. Here we used a Syrian hamster model to assess the antiviral efficacy of favipiravir, understand its mechanism of action and determine its pharmacokinetics. When treatment was initiated before or simultaneously to infection, favipiravir had a strong dose effect, leading to dramatic reduction of infectious titers in lungs and clinical alleviation of the disease. Antiviral effect of favipiravir correlated with incorporation of a large number of mutations into viral genomes and decrease of viral infectivity. The antiviral efficacy observed in this study was achieved with plasma drug exposure comparable with those previously found during human clinical trials and was associated with weight losses in animals. Thereby, pharmacokinetic and tolerance studies are required to determine whether similar effects can be safely achieved in humans.

## Introduction

In March 2020, the World Health Organization declared coronavirus disease 2019 (COVID-19) a pandemic^1^. The COVID-19 outbreak was originally identified in Wuhan, China, in December 2019 and spread rapidly around the world within a few months. The severe acute respiratory syndrome coronavirus 2 (SARS-CoV-2), the causative agent of COVID-19, belongs to the *Coronaviridae* family and is closely related to the SARS-CoV which emerged in China in 2002^2^. After an incubation period of about 5 days, disease onset usually begins with an influenza-like syndrome associated with high virus replication in respiratory tracts^3,4^. In some patients, a late acute respiratory distress syndrome, associated with high levels of inflammatory proteins, occurs within one to two weeks^3^. As of 7 July 2020, more than 11.6 million cases of COVID-19 have resulted in more than 538,000 deaths^5^. In the face of this ongoing pandemic and its unprecedented repercussions, not only on human health but also on society, ecology and economy, there is an urgent need for effective infection prevention and control measures.

Whilst host-directed and immune-based therapies could prove useful for the clinical management of critically ill patients, the availability of safe and effective antiviral molecules would represent an important step towards fighting the current pandemic. As conventional drug development is a slow process, repurposing of drugs already approved for any indication was extensively explored and led to the implementation of many clinical trials for the treatment of COVID-19^6^. However, the development of effective antiviral drugs for the treatment of COVID-19, should, as much as possible, rely on robust pre-clinical *in vivo* data, not only on efficacy generated *in vitro*. Accordingly, rapid implementation of rodent and non-human primate animal models should help to assess more finely the potential safety and efficacy of drug candidates and to determine appropriated dose regimens in humans^7,8^.

Favipiravir (6-fluoro-3-hydroxypyrazine-2-carboxamine) is an anti-influenza drug approved in Japan that has shown broad-spectrum antiviral activity against a variety of other RNA viruses^9-15^. Favipiravir is a prodrug that is metabolized intracellularly into its active ribonucleoside 5’-triphosphate form that acts as a nucleotide analogue to selectively inhibit RNA-dependent RNA polymerase and induce lethal mutagenesis^16,17^. Recently, several studies reported *in vitro* inhibitory activity of favipiravir against SARS-CoV-2 with 50% effective concentrations (EC_50_) ranging from 62 to >500µM (10 to >78µg/mL)^18-20^. Based on these results, more than 20 clinical trials on the management of COVID-19 by favipiravir are ongoing (https://clinicaltrials.gov/). In the present study, a Syrian hamster model (*Mesocricetus auratus*) was implemented to explore the *in vivo* safety and efficacy and the pharmacokinetics (PK) of several dosing regimens of favipiravir.

## Results

### *In vitro* efficacy of favipiravir

Using VeroE6 cells and an antiviral assay based on reduction of cytopathic effect (CPE), we recorded EC_50_ and EC_90_ of 32 and 52.5 µg/mL using a multiplicity of infection (MOI) of 0.001, 70.0 and >78.5µg/mL with an MOI of 0.01 (Figure S1) in accordance with previous studies^18-20^. Infectious titer reductions (fold change in comparison with untreated cells) were ≥2 with 19.6µg/mL of favipiravir and ranged between 11 and 342 with 78.5µg/mL. Using CaCo2 cells, which do not exhibit CPE with SARS-CoV-2 BavPat1 strain, infectious titer reductions were around 5 with 19.6µg/mL of favipiravir and ranged between 144 and 7721 with 78.5µg/mL of the drug. 50% cytotoxic concentrations (CC_50_) in VeroE6 and CaCo2 cells were >78.5µg/mL.

### Infection of Syrian hamsters with SARS-CoV-2

Following Chan *et al*., we implemented a hamster model to study the efficacy of antiviral compounds^7^. Firstly, we intranasally infected four-week-old female Syrian hamsters with 10^6^ TCID_50_ of virus. Groups of two animals were sacrificed 2, 3, 4 and 7 days post-infection (dpi). Viral replication was quantified in sacrificed animals by RT-qPCR in organs (lungs, brain, liver, small/large bowel, kidney, spleen and heart) and plasma. Viral loads in lungs peaked at 2 dpi, remained elevated until 4 dpi and dramatically decreased at 7 dpi (Figure 1a). Viral loads in plasma peaked at 3 dpi and viral replication was detected in the large bowel at 2 dpi (Figure 1b and Table S1). No viral RNA was detected in almost all the other samples tested (Table S1). Subsequently, we infected animals with two lower doses of virus (10^5^ and 10^4^ TCID_50_). Viral RNA was quantified in lungs, large bowel and plasma from sacrificed animals 2, 3, 4 and 7 dpi (Figure 1a and 1b). Viral loads in lungs peaked at 2 and 3 dpi with doses of 10^5^ and 10^4^ TCID_50_ respectively. Maximum viral loads in lungs of animals infected with each dose of virus were comparable. Viral RNA yields in plasma and large bowel followed a similar trend but with more variability, with this two lower doses. In addition, clinical monitoring of animals showed no marked symptoms of infection but significant weight losses from 3 dpi when compared to animals intranasally inoculated with sodium chloride 0.9% (Figure 1c).

**Figure 1:**
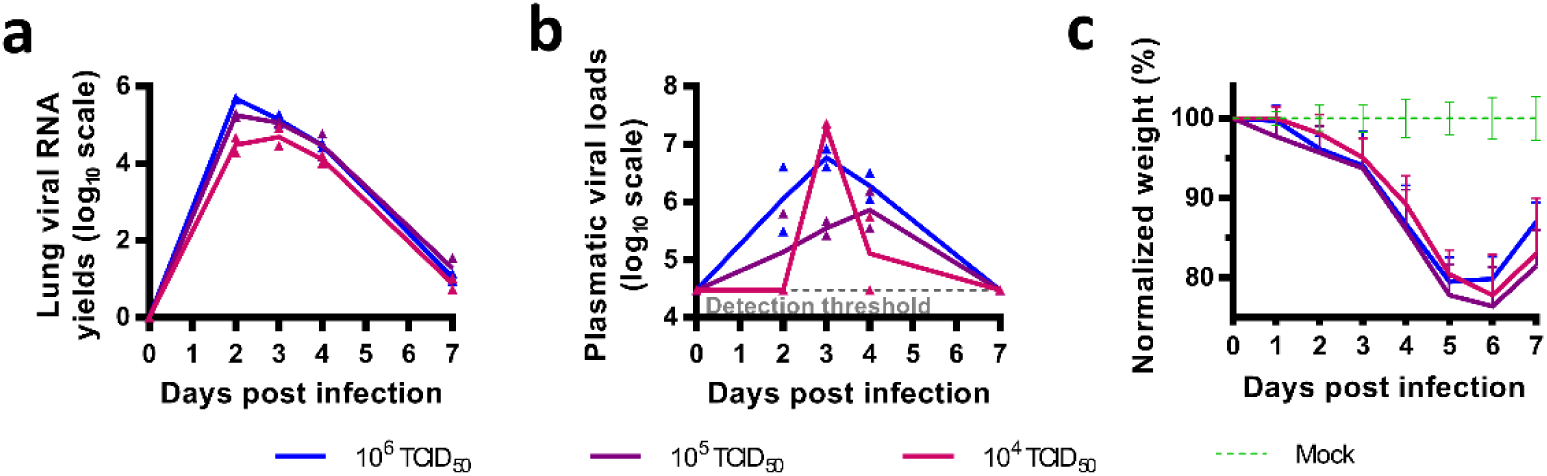
Implementation of hamster model. Hamsters were intranasally infected with 10^6^, 10^5^ or 10^4^ TCID_50_ of virus. Viral replication was quantified using an RT-qPCR assay. **a** Lung viral RNA yields. **b** Plasmatic viral loads. **c** Clinical course of the disease. Normalized weight at day *n* was calculated as follows: (% of initial weight of the animal at day *n*)/(mean % of initial weight for mock-infected animals at day *n*). Data represent mean ±SD (details in Table S1).

### *In vivo* efficacy of favipiravir

To assess the efficacy of favipiravir, hamsters received the drug, intraperitoneally, three times a day (TID). We used three doses of favipiravir: 18.75, 37.5 and 75mg/day (corresponding to 340±36, 670±42 and 1390±126 mg/kg/day respectively).

In a first set of experiments, treatment was initiated at day of infection (preemptive antiviral therapy) and ended at 2 dpi. We infected groups of 6 animals intranasally with three doses of virus (10^6^, 10^5^ and 10^4^ TCID_50_) and viral replication was measured in lungs and plasma at 3 dpi (Figure 2a). When analysis of virus replication in clarified lung homogenates was based on infectious titers (as measured using TCID_50_ assay), the effect of favipiravir in reducing infectious titers was dose dependent, in particular when low doses of virus were used to infect animals (Figure 2b). At each dose of virus, mean infectious titers for groups of animals treated with 75mg/day TID were significantly lower than those observed with untreated groups (*p*≤0.0001): reduction of infectious titers ranged between 1.9 and 3.7 log_10_. For animals infected with 10^5^ or 10^4^ TCID_50_, significant infectious titer reductions of around 0.8 log_10_ were also observed with the dose of 37,5mg/day TID (*p*≤0.038). Drug 90% and 99% effective doses (ED_90_ and ED_99_) were estimated based on these results and ranged between 31-42mg/day and 53-70mg/day respectively (Table 2). When analysis of virus replication in clarified lung homogenates were assessed on viral RNA yields (as measured using quantitative real time RT-PCR assay), significant differences with groups of untreated animals, ranging between 0.7 and 2.5 log_10_, were observed only with the higher dose of favipiravir (*p*≤0.012). Once again, this difference was more noticeable with lower doses of virus (Figure 2b). Since we found higher reductions of infectious titers than those observed with viral RNA yields, we estimated the relative infectivity of viral particle (*i*.*e*. the ratio of the number of infectious particles over the number of viral RNA molecules). Decreased infectivity was observed in all treated groups of animals. These differences were always significant with the higher dose of favipiravir (*p*≤0.031) and were significant with the dose of 37.5mg/day TID for animals infected with 10^5^ or 10^4^ TCID_50_ of virus (*p*≤0.041). We then measured plasmatic viral loads using quantitative real time RT-PCR assay and found, with the higher dose of favipiravir and the groups of animals infected with 10^6^ or 10^4^ TCID_50_, significant reductions of 2.1 and 2.62 log_10_, respectively (*p*≤0.022) (Figure 2b).

**Table 2:** Drug effective doses (ED) on reducing viral titers according to the level of viral inoculum

|  | ED<sub>50</sub> | ED<sub>90</sub> | ED<sub>99</sub> |
| --- | --- | --- | --- |
|  | mg/day (95%CI<sup>1</sup>) | mg/day (95%CI<sup>1</sup>) | mg/day (95%CI<sup>1</sup>) |
| **Preemptive therapy** |  |  |  |
| 10<sup>4</sup> TCID<sub>50</sub> | 34 (30-37) | 42 (38-46) | 53 (48-58) |
| 10<sup>5</sup> TCID<sub>50</sub> | 26 (21-30) | 37 (31-44) | 56 (46-65) |
| 10<sup>6</sup> TCID<sub>50</sub> | 15 (10-20) | 31 (21-41) | 70 (48-93) |
| **Preventive therapy** |  |  |  |
| 10<sup>4</sup> TCID<sub>50</sub> | 27 (25-29) | 35 (32-38) | 47 (44-51) |

<sup>1</sup>: 95% confidence interval
Dose-response curves are presented in Figure S2.

**Figure 2:**
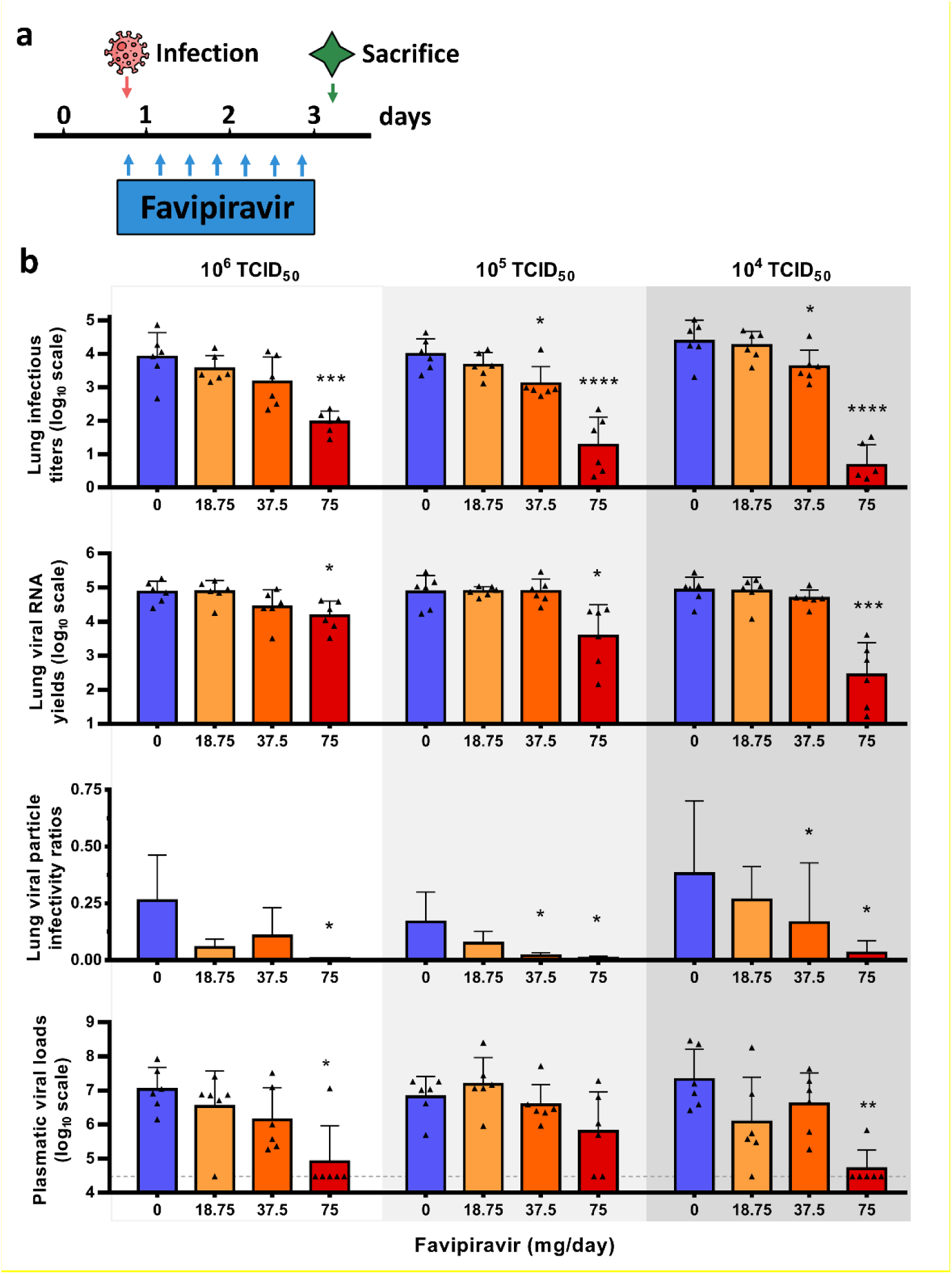
Virological results with preemptive favipiravir therapy. **a** Experimental timeline. **b** Viral replication in lungs and plasma. Hamsters were intranasally infected with 10^6^, 10^5^ or 10^4^ TCID_50_ of virus. Lung infectious titers (measured using a TCID_50_ assay) and viral RNA yields were (measured using an RT-qPCR assay) expressed in TCID_50_/copy of γ-actine gene and viral genome copies/copy of γ-actine gene respectively. Relative lung viral particle infectivities were calculated as follows: ratio of lung infectious titer over viral RNA yields. Plasmatic viral loads (measured using an RT-qPCR assay) are expressed in viral genome copies/mL of plasma (the dotted line indicates the detection threshold of the assay). Data represent mean ±SD. ****, ***, ** and * symbols indicate that the average value for the group is significantly lower than that of the untreated group with a p-value <0.0001, ranging between 0.0001-0.001, 0.001-0.01 and 0.01-0.05 respectively (details in Table S2 and S3).

In a second set of experiments, we assessed, over a period of 7 days, the impact of treatment on the clinical course of the disease using weight loss as the primary criterion (Figure 3a). Beforehand, we evaluated the toxicity of the three doses of favipiravir with groups of four non-infected animals treated from day 0 to day 3 (Figure 3b). High toxicity was observed with the dose of 75mg/day TID with significant weight loss noticed from the first day of treatment (Table S4). We also found a constant, but moderate, toxicity with the dose of 37.5mg/day TID that was significant at day 4 and 5 only. No toxicity was detected with the lower dose of favipiravir. To assess if the toxicity observed with the highest dose of favipiravir was exacerbated by the infection, we compared weight losses of infected and non-infected animals treated with the dose of 75mg/day TID. Regardless of the dose of virus, no significant difference was observed at 1, 2 and 3 dpi (Figure S3). After this evaluation of favipiravir toxicity, we intranasally infected groups of 10 animals with two doses of virus (10^5^ or 10^4^ TCID_50_). Treatment with a dose of 37.5mg/day TID was initiated on the day of infection (preemptive antiviral therapy) and ended at 3 dpi (Figure 3a). With both doses of virus, treatment was associated with clinical alleviation of the disease (Figure 3c-d). With the dose of 10^5^ TCID_50_, mean weights of treated animals were significantly higher than those of untreated animals at 5 and 6 dpi (*p*≤0.031). Similar observations were made with the dose of 10^4^ TCID_50_ at 5, 6 and 7 dpi (*p*<0.0001).

**Figure 3:**
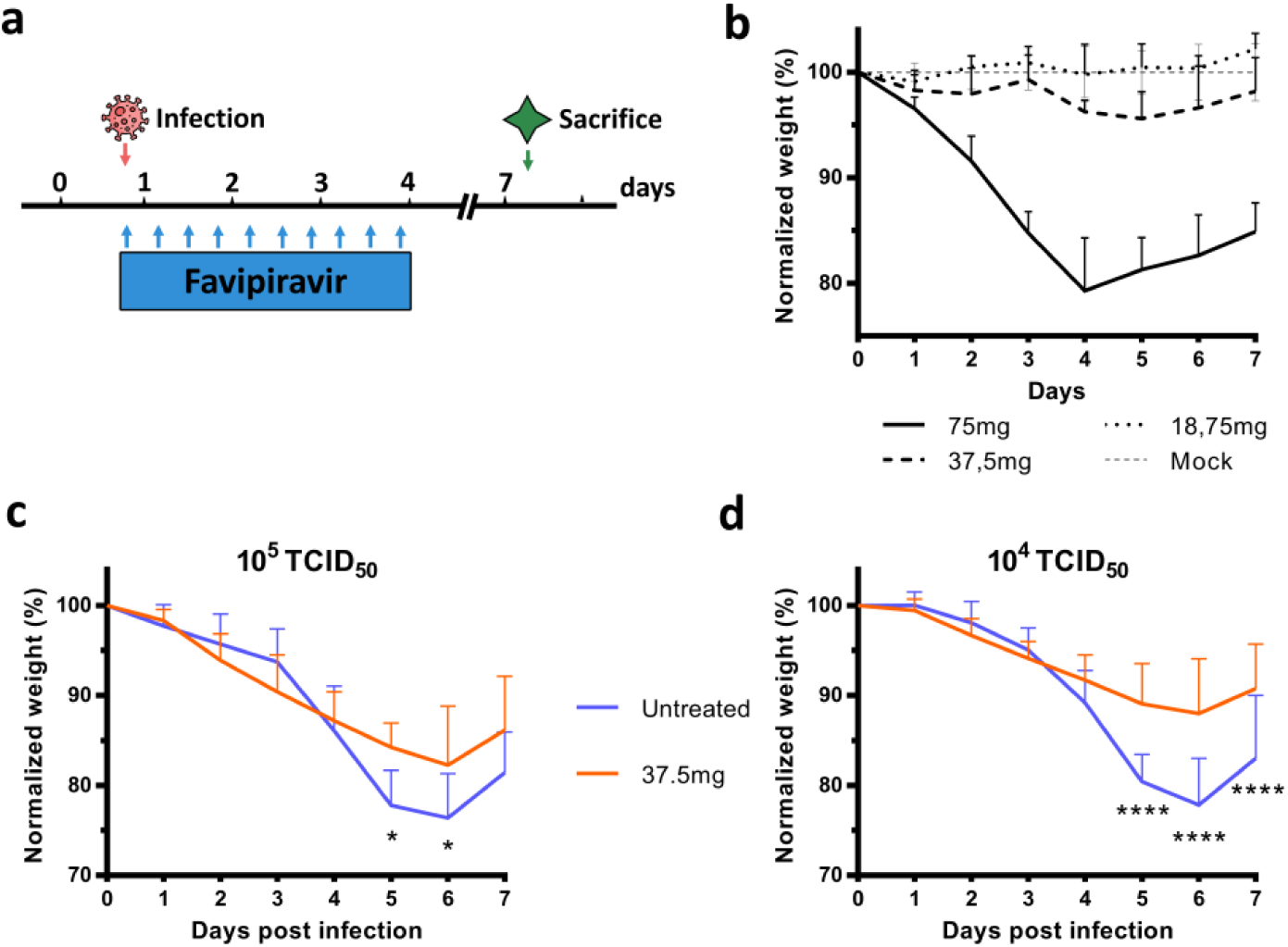
Clinical follow-up of animals. **a** Experimental timeline. **b** Evaluation of the toxicity of the three doses of favipiravir (mg/day TID) with uninfected animals following an identical experimental timeline without infection. **c-d** Clinical follow-up with animals infected respectively with 10^5^ and 10^4^ TCID_50_ of virus and treated with a dose of favipiravir of 37.5mg/day TID. Normalized weight at day *n* was calculated as follows: (% of initial weight of the animal at day *n*)/(mean % of initial weight for mock-infected animals at day *n*). Data represent mean ±SD. **** and * symbols indicate a significant difference between treated and untreated animals with a p-value <0.0001 and ranging between 0.01-0.05 respectively (details in Table S2 and S4).

In a third set of experiments, treatment was started one day before infection (preventive antiviral therapy) and ended at 2 dpi. We intranasally infected groups of 6 animals with 10^4^ TCID_50_ of virus and viral replication was measured in lungs and plasma at 3 dpi (Figure 4a). Once again, an inverse relationship was observed between lung infectious titers and the dose of favipiravir (Figure 4b). Mean infectious titers for groups of animals treated with 37.5 and 75mg/day TID were significantly lower than those observed with untreated groups (*p*≤0.002). Of note, undetectable infectious titers were found for all animals treated with the higher dose. Estimated ED_90_ and ED_99_ were 35 and 47mg/day respectively (Table 2). Significant reductions of viral RNA yields of 0.9 and 3.3 log_10_, were observed with animals treated with 37.5 and 75mg/day TID respectively (*p*≤0.023). Resulting infectivity of viral particle was decreased, with a significant reduction only for the higher dose of favipiravir (*p*=0.005). Finally, we found significantly reduced plasmatic viral loads with animals treated with 37.5 and 75mg/day TID (*p*≤0.005).

**Figure 4:**
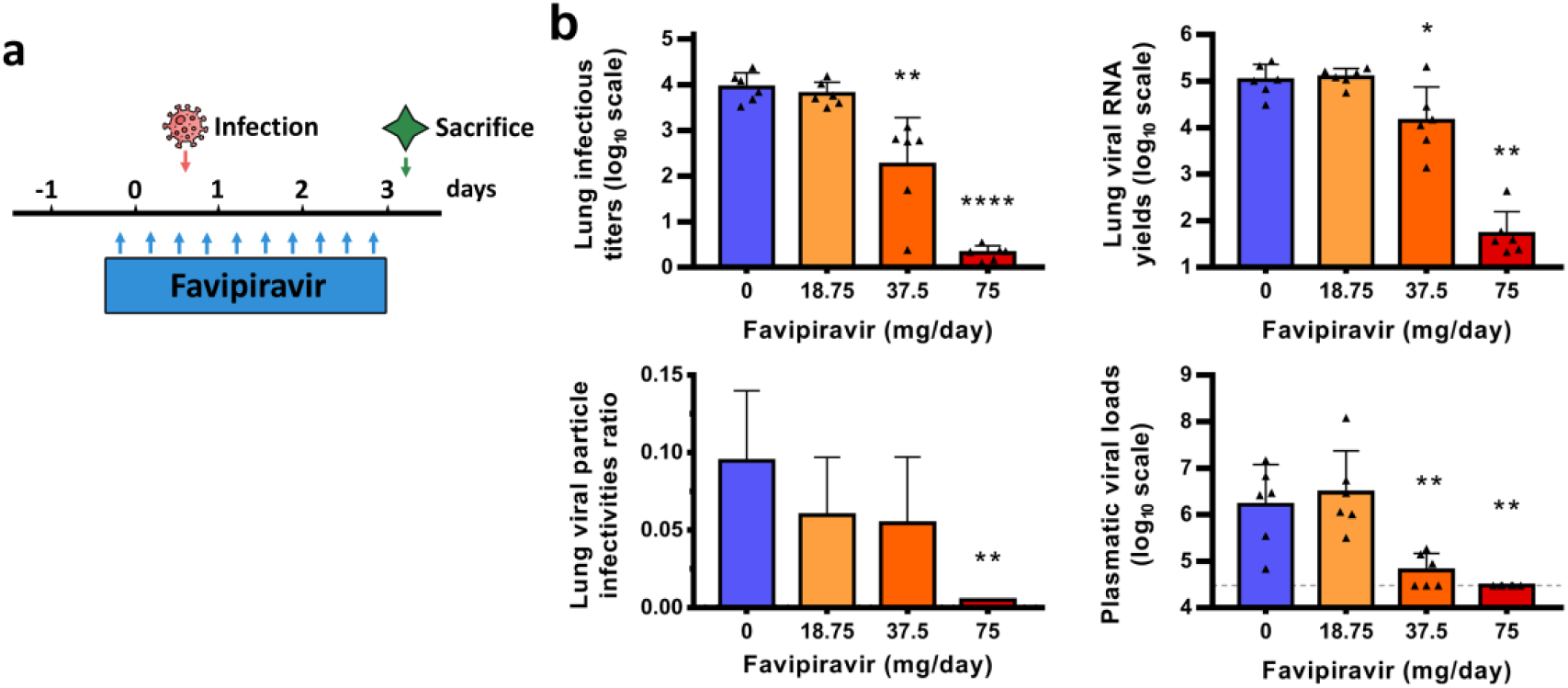
Virological results with preventive favipiravir therapy. **a** Experimental timeline. **b** Viral replication in lungs and plasma. Hamsters were intranasally infected with 10^4^ TCID_50_ of virus. Lung infectious titers (measured using a TCID_50_ assay) and viral RNA yields awee (measured using an RT-qPCR assay). They are expressed in TCID_50_/copy of γ-actine gene and viral genome copies/copy of γ-actine gene respectively. Relative lung virus infectivities were calculated as follows: ratio of lung infectious titer over viral RNA yields. Plasmatic viral loads (measured using an RT-qPCR assay) are expressed in viral genome copies/mL of plasma (the dotted line indicates the detection threshold of the assay). Data represent mean ±SD. ****, ** and * symbols indicate that the average value for the group is significantly different from that of the untreated group with a p-value <0.0001, ranging between 0.001-0.01 and 0.01-0.05 respectively (details in Table S2 and S3).

### Favipiravir pharmacokinetics (PK) in a hamster model

We first assessed the PK and lung distribution of favipiravir in a subgroup of uninfected animals. Groups of animals were treated respectively with a single dose of favipiravir administrated intraperitoneally: 6.25mg, 12.5mg and 25mg. In each dose group, we sacrificed 3 animals at specific time points post-treatment (0.5, 1, 5 or 8 hours) for determination of favipiravir in plasma. Drug concentration in lung tissue was determined at 0.5 and 5 hours post-treatment. Subsequently, we assessed the favipiravir concentration after multiple dose in animals intranasally infected with 10^5^ TCID_50_ of virus. Groups of 9 animals received the three doses evaluated for 3 days (Figure 2a): 18.75mg/day, 37.5mg/day and 75mg/day TID and were sacrificed at 12-hours after the last treatment dose. Favipiravir was quantified in plasma (n=9) and lung tissue (n=3).

Results are presented in Table 3 and Figure S4. The single dose PK analysis showed that the maximum concentration of favipiravir was observed at 0.5 hour at all doses, then plasma drug concentrations decreased exponentially to reach concentrations below 10µg/ml at 12 hours. Favipiravir PK exhibited a non-linear increase in concentration between the doses. After multiple doses, trough concentrations (12 hours) of favipiravir also exhibited a non-linear increase between doses. The extrapolated 12 hours post-treatment concentrations after a single dose were calculated in order to determine the accumulation ratio. Accumulation ratios were respectively 6, 16 and 21 at the 3 doses, confirming the non-proportional increase between doses. The average concentration after single dose administration over 0 to 12-hour intervals was calculated and the respective values obtained were 10.1µg/mL, 38.7µg/mL and 100.5µg/mL for the 3 favipiravir doses.

**Table 3:**
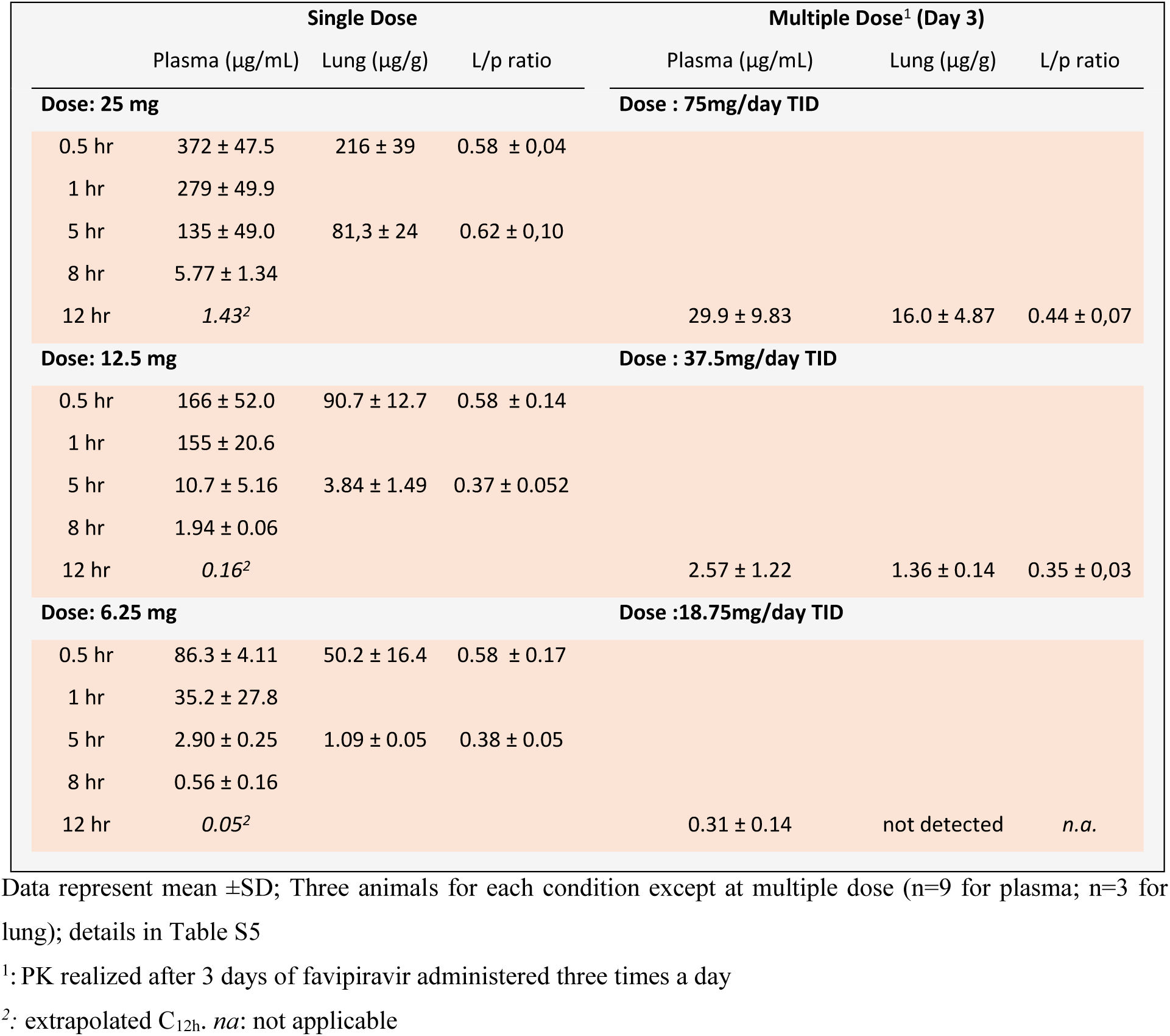
Plasma and lung concentrations of favipiravir after administration of a single dose or multiple dose of favipiravir

Favipiravir lung concentrations were 1.6 to 2.7-fold lower than in plasma for both administration of single and multiple doses. After a single dose, the mean lung to plasma ratio ranged from 0.37 to 0.62 according to the time post-treatment and was similar between the 3 doses of favipiravir at 0.5 hours. A high ratio 5 hours post-treatment was observed at the highest dose (25mg) with an increase by a factor 1.6 to 1.8 compared with the lower doses. After multiple doses, the lung penetration of favipiravir was confirmed with a mean lung to plasma ratio ranging from 0.35 to 0.44. Favipiravir was not detected in the lungs at the lowest dose (18.75mg/day).

### Mutagenic effect of favipiravir

To understand which genomic modifications accompanied favipiravir treatment, direct complete genome sequencing of clarified lung homogenates from animals intranasally infected with 10^6^ TCID_50_ of virus and treated with the two highest doses of drug (preemptive antiviral therapy; Figure 2) was performed. Data were generated by next generation sequencing from lung samples of four animals per group (untreated, 37.5mg/day TID and 75mg/day TID). The mean sequencing coverage for each sample ranged from 10,991 to 37,991 reads per genomic position and we subjected substitutions with a frequency ≥1% to further analysis. The genetic variability in virus stock was also analyzed: 14 nucleotide polymorphisms were detected of which 5 recorded a mutation frequency higher than 10% (Table S6).

In order to study the mutagenic effect of favipiravir, we used the consensus sequence from virus stock as reference and all the mutations simultaneously detected in a lung sample and in virus stock were not considered in the further analysis (1 to 4 mutations per sample, see Table S6). Overall, no majority mutations were detected (mutation frequency >50%), mutations were distributed throughout the whole genome and almost all of them exhibited a frequency lower than 10% (Figure 5a and 5b).

**Figure 5:**
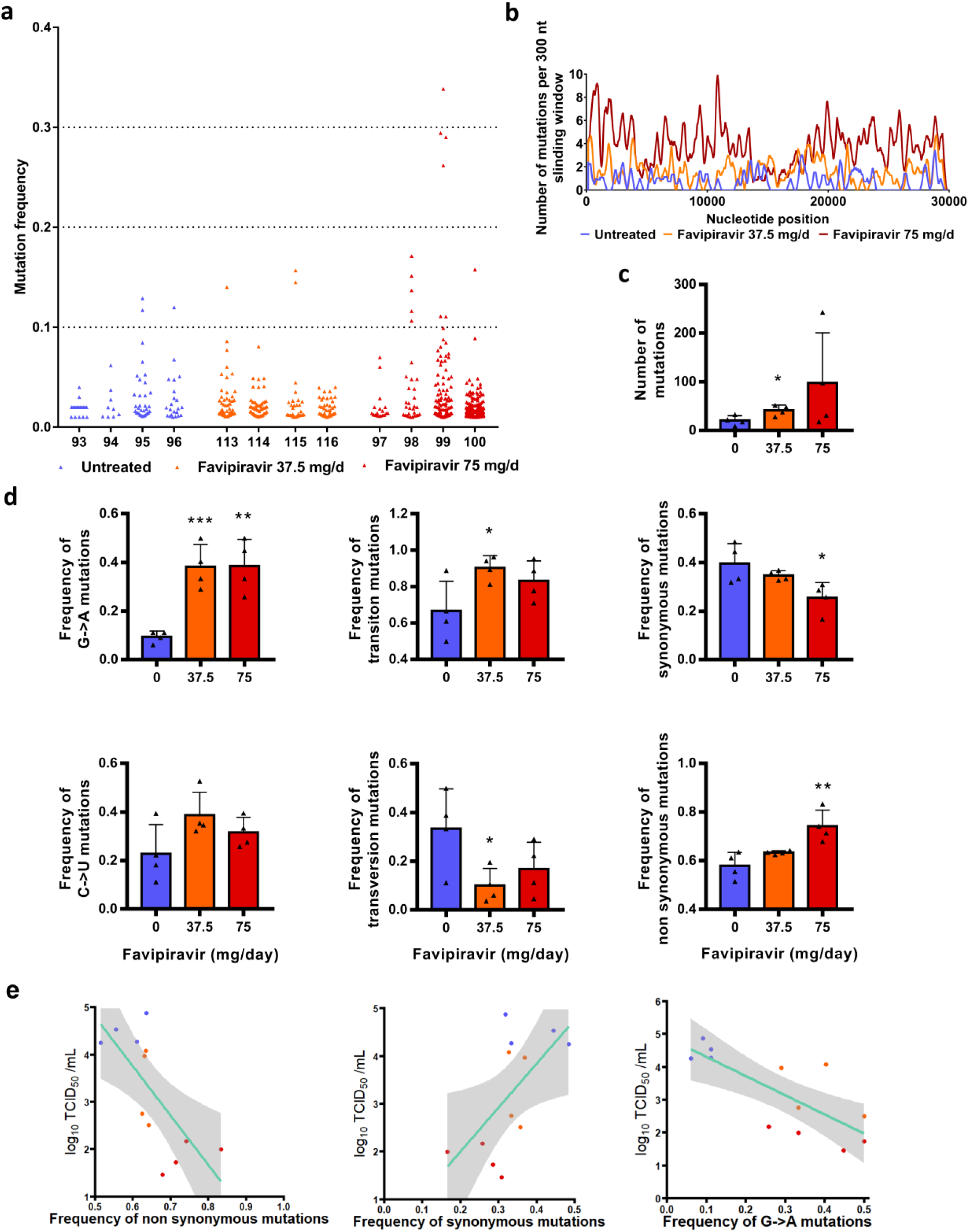
Mutagenic effect of favipiravir. **a** Viral genetic diversity in clarified lung homogenates. For each condition, four samples were analyzed. Each triangle represents a mutation (only substitutions with a frequency ≥1% were considered). **b** Patterns of mutation distribution on complete viral genome. Each variable nucleotide position was counted only once when found. The variability was represented using 75 nt sliding windows. For each condition, variable nucleotide positions were determined and represented using a 300 nt sliding window. **c** Mean number of mutations. Data represent mean ±SD. **d** Mutation characteristics. For each sample, the frequency of a given mutation was calculated as follows: number of this kind of mutation detected in the sample divided by the total number of mutations detected in this sample. Data represent mean ±SD. ** and * symbols indicate that the average value for the group is significantly different from that of the untreated group with a p-value ranging between 0.001-0.01 and 0.01-0.05 respectively (details in details in Table S6 and S7). **e** Association between lung infectious titers (measured using a TCID_50_ assay) and frequency of non synonymous, synonymous and G→A mutations. Each dot represent data from a given animal.

Results revealed a relationship between the number of mutations detected per sample and the dose of favipiravir (Figure 5c): the mean number of mutations increased by a factor 2 and 4.8 with groups of animals treated with 37.5 and 75mg/day TID, respectively. The difference is significant only with a dose of 37.5mg/day TID (*p*=0.029). This increase of the number of mutations is mainly the consequence of the occurrence of a large number of G→A substitutions and, to a lesser extent, C→U substitutions. Consequently, regardless of the dose of favipiravir, mean frequency of G→A substitutions was significantly increased by a factor of 4.2 (p≤0.009). This rise of these transition mutations led to increased frequency of all transition mutations (significant only at dose of 37.5mg/day TID; p=0.037) and increased frequency of non-synonymous mutations (significant only at dose of 75mg/day TID; p=0.009) (Figure 5d). We investigated whether or not effectiveness in treated animals was linked with the characteristics of the mutations detected on viral populations and found that infectious titers in lungs were negatively associated with frequency of non-synonymous and G→A mutations, and positively associated with frequency of synonymous mutations (p<0.03; Figure 5e). Finally, our experiments revealed some parallel evolution events; 32 substitutions in viral sub-populations were detected in two independent animals. Notably, 18 of these shared mutations were detected only with treated animals, 14 of them being non-synonymous (Table S8). These mutations are located in nsp2, 3, 4, 5, 6, 14, N protein, Matrix, ORF 3a and 8. At this stage, one cannot conclude if these substitutions reflect the adaptation to the hamster model or are the result of the antiviral selection.

## Discussion

In the current study, we used a hamster model to assess efficacy of the favipiravir against SARS-CoV-2. Following infection, viral RNA was mainly detected in lungs, blood, and, to a lesser extent, in the large bowel. Peak of viral replication was observed at 2-3 dpi followed by observation of significant weight losses, in line with recently reported investigations that involved 6-10 weeks old hamsters^7,21^. Clinically, the main symptom was weight loss, observed from the first day of infection and followed by recovery at 6dpi. This confirmed that the *in vivo* model, with younger animals (4 weeks-old), is suitable for preclinical evaluation of antiviral compounds against SARS-CoV-2.

Using a preemptive strategy, we demonstrated that doses of favipiravir of around 700-1400mg/kg/day TID reduced viral replication in lungs of infected animals and allowed clinical alleviation of the disease. In the most favourable situation, where high doses were used as a preventive therapy, favipiravir led to undetectable viral replication in lung and plasma. These results showed that the use of high doses of favipiravir could expand its *in vivo* spectrum against RNA viruses.

Reduction of viral replication was greater when estimated on the basis of infectious titers than on total viral RNA as previously observed in non-human primates treated with Remdesivir^22^. However, the effective doses of favipiravir were higher than those usually used in rodent models (≈100-400mg/kg/day)^10,12,23-26^. This can be correlated with the high favipiravir EC_50_ found *in vitro* for SARS-CoV-2. Moreover, effective doses were associated with significant toxicity in our hamster model. This observed toxicity reflected only the adverse effects of favipiravir and was not exacerbated during SARS-CoV-2 infection. Indeed, similar weight losses were measured among infected and non-infected animals treated with the highest dose of favipiravir at 1, 2 and 3dpi.

In the present study, reduction of viral replication was correlated with the dose of favipiravir administrated and inversely correlated with the dose of virus inoculated. In a recent study, favipiravir administrated *per os* twice daily (loading dose of 600mg/kg/day followed by 300mg/kg/day) revealed a mild reduction of lung viral RNA yields using a similar hamster model with high doses of virus (2×10^6^ TCID_50_)^21^. These results are in accordance with ours at the lower dose of favipiravir (around 340mg/kg/day TID).

With influenza viruses, favipiravir acts as a nucleotide analogue. It is metabolized intracellularly to its active form and incorporated into nascent viral RNA strands. This inhibits RNA strand extension and induces abnormal levels of mutation accumulation into the viral genome^16,17^. Recently, it was shown *in vitro* that favipiravir has a similar mechanism of action with SARS-CoV-2 through a combination of chain termination, reduced RNA synthesis and lethal mutagenesis^20^. Our genomic analysis confirmed the mutagenic effect of favipiravir *in vivo*. Indeed, we found that favipiravir treatment induced appearance of a large number of G→A and C→U mutations into viral genomes. This was associated to a decrease of viral infectivity probably because alteration of the genomic RNA disturb the replication capacity. Similar findings were described *in vitro* and *in vivo* with other RNA viruses^9,16,27,28^. Of note, we also observed a strong inverse association between infectious titers in lungs and the proportion of non-synonymous mutations detected in viral populations. Because random non-synonymous mutations are more deleterious than synonymous mutations^29^, this suggests that they were randomly distributed over the three positions of the codons and that no compensatory mechanism was triggered by the virus to eliminate them (*i*.*e*. negative selection). Finally, the inverse correlation between lung infections titers and the frequency of G→A substitutions showed that an increased proportion of these mutations beyond an error threshold might be expected to cause lethal mutagenesis.

Genomic analyses revealed that 18 mutations detected in viral sub-populations were shared only with treated animals. Two of them were located in the nsp14 coding region involved in the proof-reading activity of the viral RNA polymerisation^30,31^. However, they were located in the N7 MTase domain involved in viral RNA capping^32,33^. By comparison, resistance mutations selected against Remdesivir in β-coronavirus murine hepatitis virus model were obtained in the RdRP (nsp12) coding sequence^34^. Further investigations are needed to assess the impact of these mutations on the antiviral effect of favipiravir.

Favipiravir PK in our hamster model displayed a non-linear increase in plasma exposure between the doses as already reported in nonhuman primates^35^. The observed favipiravir concentration versus time profiles were in agreement with previous results of a PK study performed in 7-8 week-old hamsters orally treated with a single dose of 100mg/kg of favipiravir^36^. The maximum plasma drug concentration occurred at 0.5 h after oral administration, earlier than in humans, and then decreased rapidly in agreement with its short half-life^37^. After repeated doses, plasma exposure confirmed non-linear PK over the entire range of doses, further emphasized by accumulation ratios. The important accumulation observed at the highest dose could explain in part the toxicity observed in hamsters at this dose. Favipiravir undergoes an important hepatic metabolism mainly by aldehyde oxidase producing an inactive M1 metabolite and inhibits aldehyde oxidase activity in a concentration- and time-dependent manner. These properties explain the self-inhibition of its own metabolism as observed in our study in which the highest dose of favipiravir led to a greater increase in favipiravir concentrations^38^.

A good penetration of favipiravir in lungs was observed with lung/plasma ratios ranging from 35 to 44% after repeated doses, consistent with its physicochemical properties. Lung exposure was also in accordance with previous studies^36^.

The medium dose of favipiravir used in this study (670mg/kg/day TID) is within the range of the estimated doses required to reduce by 90% (ED90) the level of infectious titers in lungs (ranging between 570 and 780mg/kg/day). Animals treated with this dose displayed significant reduction of viral replication in lungs, limited drug-associated toxicity and clinical alleviation of the disease. Regarding the accumulation ratio after repeated doses and the good penetration of favipiravir in lungs, effective concentrations can be expected in lungs, throughout the course of treatment using this dose of 670mg/kg/day TID.

How clinically realistic are these results? To address this question we compared the drug concentrations obtained in the hamster model with those obtained in patients. In 2016, a clinical trial evaluated the use of favipiravir in Ebola infected patients^39^. The dose used in Ebola infected patients was 6000mg on day 0 followed by 1200mg BID for 9 days. The median trough concentrations of favipiravir at Day 2 and Day 4 were 46.1 and 25.9µg/mL, respectively. This is within the range observed here in hamsters treated with the highest dose (around 1400mg/kg/day), with a mean trough concentration of 29.9µg/mL. However, additional investigations are required to determine whether or not similar favipiravir plasma exposure in SARS-COV-2 infected patients are associated with antiviral activity. The major differences in PK between hamster and humans, and the toxicity observed at the highest doses in our animal model limits the extrapolation of our results. Therefore, whether safe dosing regimens in humans may achieve similar plasma exposure and recapitulate the profound effect on viral replication is unknown. Further, the intracellular concentration of the active metabolite was not determined and which parameter of the drug pharmacokinetics best drives the antiviral effect remains to be established.

In summary, this study establishes that high doses of favipiravir are associated with antiviral activity against SARS-CoV-2 infection in a hamster model. The better antiviral efficacy was observed using a preventive strategy, suggesting that favipiravir could be more appropriate for a prophylactic use. Our results should be interpreted with caution because high doses of favipiravir were associated with signs of toxicity in our model. It is required to determine if a tolerable dosing regimen could generate similar exposure in non-human primates, associated with significant antiviral activity, before testing a high dose regimen in COVID-19 patients. Furthermore, subsequent studies should determine if an increased antiviral efficacy can be reached using favipiravir in association with other effective antiviral drugs, since this strategy may enable to reduce the dosing regimen of favipiravir. Finally, this work reinforces the need for rapid development of animal models to confirm *in vivo* efficacy of antiviral compounds and accordingly, to determine appropriate dose regimens in humans before starting clinical trials.

## Methods

### Cells

VeroE6 cells (ATCC CRL-1586) and Caco-2 cells (ATCC HTB-37) were grown at 37°C with 5% CO_2_ in minimal essential medium (MEM) supplemented with 7.5% heat-inactivated fetal bovine serum (FBS), 1% Penicillin/Streptomycin and 1% non-essential amino acids (all from ThermoFisher Scientific).

### Virus

All experiments with infectious virus were conducted in biosafety level (BSL) 3 laboratory. SARS-CoV-2 strain BavPat1, supplied through European Virus Archive GLOBAL (https://www.european-virus-archive.com/), was provided by Christian Drosten (Berlin, Germany). Virus stocks were prepared by inoculating at MOI of 0.001 a 25cm2 culture flask of confluent VeroE6 cells with MEM medium supplemented with 2.5% FBS. The cell supernatant medium was replaced each 24h hours and harvested at the peak of infection, supplemented with 25mM HEPES (Sigma), aliquoted and stored at −80°C.

### *In vitro* determination of EC_50_, EC_90,_ CC_50_ and infectious titer reductions

One day prior to infection, 5×10^4^ VeroE6 cells were seeded in 96-well culture plates (5×10^4^ cells/well in 100µL of 2.5% FBS medium (assay medium). The next day, seven 2-fold serial dilutions of favipiravir (Courtesy of Toyama-Chemical; 0.61µg/mL to 78.5µg/mL, in triplicate) were added (25µL/well, in assay medium). Eight virus control wells were supplemented with 25µL of assay medium and eight cell controls were supplemented with 50µL of assay medium. After 15 min, 25µL of virus suspension, diluted in assay medium, was added to the wells at an MOI of 0.01 or 0.001 (except for cell controls). Three days after infection, cell supernatant media were collected to perform TCID_50_ assay (at concentration of 78.5, 39.3, 19.6µg/mL), as described below, in order to calculate infectious titer reductions and cell viability was assessed using CellTiter-Blue reagent (Promega) following manufacturer’s intructions. Fluorescence (560/590nm) was recorded with a Tecan Infinite 200Pro machine (Tecan). The 50% and 90% effective concentrations (EC_50_, EC_90_) were determined using logarithmic interpolation (% of inhibition were calculated as follows: (OD_sample_-OD_virus control_)/(OD_cell control_-OD_virus control_)). For the evaluation of CC_50_ (the concentration that induced 50% cytoxicity), the same culture conditions were set as for the determination of the EC_50_, without addition of the virus, then cell viability was measured using CellTiter Blue (Promega). CC_50_ was determined using logarithmic interpolation.

### *In vivo* experiments

#### Approval and authorization

*In vivo* experiments were approved by the local ethical committee (C2EA—14) and the French ‘Ministère de l’Enseignement Supérieur, de la Recherche et de l’Innovation’ (APAFIS#23975) and performed in accordance with the French national guidelines and the European legislation covering the use of animals for scientific purposes. All experiments were conducted in BSL 3 laboratory.

#### Animal handling

Three-week-old female Syrian hamsters were provided by Janvier Labs. Animals were maintained in ISOcage P - Bioexclusion System (Techniplast) with unlimited access to water/food and 14h/10h light/dark cycle. Animals were weighed and monitored daily for the duration of the study to detect the appearance of any clinical signs of illness/suffering. Virus inoculation was performed under general anesthesia (isoflurane). Organs and blood were collected after euthanasia (cervical dislocation) which was also realized under general anesthesia (isofluorane).

#### Hamster Infection

Anesthetized animals (four-week-old) were intranasally infected with 50µL containing 10^6^, 10^5^ or 10^4^ TCID_50_ of virus in 0.9% sodium chloride solution). The mock group was intranasally inoculated with 50µL of 0.9% sodium chloride solution.

#### Favipiravir administration

Hamster were intra-peritoneally inoculated with different doses of favipiravir. Control group were intra-peritoneally inoculated with a 0.9% sodium chloride solution.

#### Organ collection

Organs were first washed in 10mL of 0.9% sodium chloride solution and then transferred to a 2mL or 50mL tube containing respectively 1mL (small/large bowel pieces, kidney, spleen and heart) or 10mL (lungs, brain and liver) of 0.9% sodium chloride solution and 3mm glass beads. They were crushed using a the Tissue Lyser machine (Retsch MM400) for 5min at 30 cycles/s and then centrifuged 5min à 1200g. Supernatant media were transferred to a 2mL tube, centrifuged 10 min at 16,200g and stored at −80°C. One milliliter of blood was harvested in a 2mL tube containing 100µL of 0.5M EDTA (ThermoFischer Scientific). Blood was centrifuged for 10 min at 16,200g and stored at −80°C.

### Quantitative real-time RT-PCR (RT-qPCR) assays

To avoid contamination, all experiments were conducted in a molecular biology laboratory that is specifically designed for clinical diagnosis using molecular techniques, and which includes separate laboratories dedicated to perform each step of the procedure. Prior to PCR amplification, RNA extraction was performed using the QIAamp 96 DNA kit and the Qiacube HT kit and the Qiacube HT (both from Qiagen) following the manufacturer’s instructions. Shortly, 100 µl of organ clarified homogenates, spiked with 10µL of internal control (bacteriophage MS2)^40^, were transferred into an S-block containing the recommended volumes of VXL, proteinase K and RNA carrier. RT-qPCR (SARS-CoV-2 and MS2 viral genome detection) were performed with the Express one step RT-qPCR Universal kit (ThermoFisher Scientific) using 3.5µL of RNA and 6.5µL of RT-qPCR mix that contains 250nmol of each primer and 75nmol of probe. Amplification was performed with the QuantStudio 12K Flex Real-Time PCR System (ThermoFisher Scientific) using the following conditions: 50°C for 10min, 95°C for 20s, followed by 40 cycles of 95°C for 3s, 60°C for 30s. qPCR (γ-actine gene detection) was perfomed under the same condition as RT-qPCR with the following modifications: we used the Express one step qPCR Universal kit (ThermoFisher Scientific) and the 50°C step of the amplification cycle was removed. Primers and probes sequences used to detect SARS-CoV-2, MS2 and γ-actine are described in Table S9.

### Tissue-culture infectious dose 50 (TCID_50_) assay

To determine infectious titers, 96-well culture plates containing confluent VeroE6 cells were inoculated with 150μL per well of serial dilutions of each sample (four-fold or ten-fold dilutions when analyzing lung clarified homogenates or cell supernatant media respectively). Each dilution was performed in sextuplicate. Plates were incubated for 4 days and then read for the absence or presence of cytopathic effect in each well. Infectious titers were estimated using the method described by Reed & Muench^41^.

### Favipiravir pharmacokinetics

Animal handling, hamster infections and favipiravir administrations were performed as described above. A piece of left lung was first washed in 10mL of sodium chloride 0.9% solution, blotted with filter paper, weighed and then transferred to a 2mL tube containing 1mL of 0.9% sodium chloride solution and 3mm glass beads. It was crushed using the Tissue Lyser machine (Retsch MM400) during 10min at 30 cycles/s and then centrifuged 5min à 1200g. Supernatant media were transferred to 2mL tubes, centrifuged 10 min at 16,200g and stored at −80°C. One milliliter of blood was harvested in a 2mL tube containing 100µL of 0.5M EDTA (ThermoFischer Scientific). Blood was centrifuged for 10 min at 16,200g and stored at −80°C.

Quantification of favipiravir in plasma and lung tissues was performed by a validated sensitive and selective validated high-performance liquid chromatography coupled with tandem mass spectrometry method (UPLC-TQD, Waters, USA) with a lower limit of quantification of 0.1 µg/mL. Precision and accuracy of the 3 quality control samples (QCs) were within 15% over the calibration range (0.5 µg/mL to 100 µg/mL) (Bekegnran *et al*., submitted). Favipiravir was extracted by a simple protein precipitation method, using acetonitrile for plasma and ice-cold acetonitrile for clarified lung homogenates. Briefly, 50 µL of samples matrix was added to 500µL of acetonitrile solution containing the internal standard (favipiravir-13C,15N, Alsachim), then vortexed for 2min followed by centrifugation for 10min at 4°C. The supernatant medium was evaporated and the dry residues were then transferred to 96-well plates and 50 µL was injected. To assess the selectivity and specificity of the method and matrix effect, blank plasma and tissues homogenates from 2 control animals (uninfected and untreated) were processed at each run. Moreover, the same control samples spiked with favipiravir concentration equivalent to the QCs (0.75, 50 and 80 µg/mL) were also processed and compared to the QCs samples.

Noncompartemental analysis conducted using software Pkanalix2019R2 (www.lixoft.com). Areas under the plasma concentration time curve were computed using medians of favipiravir concentrations at 0.5, 1, 5 and 8 hours, and extrapolated until T=12h. C_trough_ were extrapolated at T=12h using lambda-z loglinear regression on the decreasing slope of concentrations.

### Sequence analysis of the full-length genome

200µL of lung clarified homogenate or infectious cell supernatant (virus stock) was inactivated with an equal volume of VXL lysis buffer (Qiagen) and viral RNA was extracted using an EZ1 Advanced XL robot with the EZ1 mini virus 2.0 kit (both from Qiagen) and linear acrylamide (ThermoFisher Scientific) in place of carrier RNA. cDNA was generated in a final volume of 40µL using 14µL of nucleic acid extract, random hexamer and the Protoscript II First Strand cDNA Synthesis Kit (New England Biolabs). A specific set of primers (Table S10) was used to generate thirteen amplicons covering the entire genome with the Q5 High-Fidelity DNA polymerase (New England Biolabs). PCR mixes (final volume 25µL) contained 2.5µL of cDNA, 2µL of each primer (10µM) and 12.5 µL of Q5 High-Fidelity 2X Master Mix. Amplification was performed with the following conditions: 30 sec at 98°C, then 45 cycles of 15 sec at 98°C and 5 min à 65°C. Size of PCR products was verified by gel electrophoresis. For each sample, an equimolar pool of all amplicons was prepared and purified using Monarch PCR & DNA Cleanup Kit (New England Biolabs). After DNA quantification using Qubit dsDNA HS Assay Kit and Qubit 2.0 fluorometer (ThermoFisher Scientific), amplicons were fragmented by sonication into fragments of around 200bp long. Libraries were built by adding barcodes, for sample identification, and primers using AB Library Builder System (ThermoFisher Scientific). To pool equimolarly the barcoded samples a quantification step by real time PCR using Ion Library TaqMan Quantitation Kit (ThermoFisher Scientific) was performed. Then, emulsion PCR from pools and loading on 530 chip was performed using the automated Ion Chef instrument (ThermoFisher Scientific). Sequencing was performed using the S5 Ion torrent technology v5.12 (ThermoFisher Scientific) following manufacturer’s instructions. Consensus sequence was obtained after trimming of reads (reads with quality score <0.99, and length <100pb were removed and the 30 first and 30 last nucleotides were removed from the reads). Mapping of the reads on a reference (determine following blast of De Novo contigs) was done using CLC genomics workbench software v.20 (Qiagen). A *de novo* contig was also produced to ensure that the consensus sequence was not affected by the reference sequence. Mutation frequency for each position was calculated as the number of reads with a mutation compared to the reference divided by the total number of reads at that site. Only substitutions with a frequency of at least 1% were taken into account for the analysis (Table S6).

### ED_50_, ED_90_ and ED_99_ determination

We conducted a nonlinear regression of infectious viral load against dose, using an E_max_ model, giving 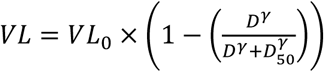 being infectious viral load of untreated animals. We estimated *D*_50_ the dose required to decrease viral load by 50%, using a coefficient *γ* to account for the high sigmoidicity of the relation between dose and titers. *γ* coefficient was chosen as the one maximizing likelihood of the model. We extrapolated the *D*_90_ and *D*_99_ using 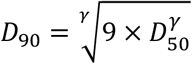 and 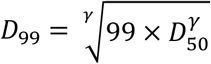, as well as their 95% confidence interval using the delta method.

### Statistical analysis

Graphical representations and statistical analyses were performed with Graphpad Prism 7 (Graphpad software) except linear/nonlinear regressions and their corresponding graphical representations that were performed using R statistical software (http://www.R-project.org). Statistical details for each experiments are described in the figure legends and in corresponding supplemental tables. P-values lower than 0.05 were considered statistically significant.

## Acknowledgments

We thank Laurence Thirion (UVE; Marseille) for providing RT-qPCR systems. We thank Camille Placidi (UVE; Marseille) for her technical contribution. We also thank Pr. Ernest A. Gould (UVE; Marseille) for his careful reading of the manuscript and English language editing. We thank Pr Drosten and Pr Drexler for providing the SARS-CoV-2 strain through the European Research infrastructure EVA GLOBAL. This work was supported by the Fondation de France “call FLASH COVID-19”, project TAMAC, by “Institut national de la santé et de la recherche médicale” through the REACTing (REsearch and ACTion targeting emerging infectious diseases) initiative (“Preuve de concept pour la production rapide de virus recombinant SARS-CoV-2”), and by European Virus Archive Global (EVA 213 GLOBAL) funded by the European Union’s Horizon 2020 research and innovation program under grant agreement No. 871029. A part of the work was done on the Aix Marseille University antivirals platform “AD2P”.

## Author Contributions

Conceptualization, J.S.D., M.C., G.M. and A.N.; Methodology, J.S.D., M.C., G.L., G.M. and A.N.; Formal Analysis, J.S.D., M.C. and G.L.; Investigation, J.S.D., M.C., G.M., F.T., P.R.P., G.P., K.B. and A.N.; Resources, F.T., B.C., J.G., X.d.L., C.S. and A.N.; Writing – Original Draft, J.S.D., M.C., J.G., C.S. and A.N.; Writing – Review & Editing, J.G., X.d.L., C.S. and A.N.; Visualization, J.S.D., M.C., G.L., F.T., P.R.P. and A.N.; Supervision, A.N.; Funding Acquisition, F.T., B.C., X.d.L. and A.N.

## Supplemental Data

Supplemental figure 1: In vitro efficacy of favipiravir

Supplemental figure 2: Dose-response curves

Supplemental figure 3: Evaluation of the toxicity for animals infected and treated with high doses of favipiravir

Supplemental figure 4: Plasma concentrations of favipiravir after administration of a single dose of favipiravir

Supplemental table 1: Implementation of hamster model

Supplemental table 2: Individual data from in vivo experiments

Supplemental table 3: Statistical analysis of in vivo experiments

Supplemental table 4: Statistical analysis of clinical monitoring

Supplemental table 5: Individual data of favipiravir pharmacokinetics

Supplemental table 6: Individual data for analysis of mutagenic effect of favipiravir

Supplemental table 7: Statistical analysis of mutagenic effect of favipiravir

Supplemental table 8: Shared mutations detected in lung clarified homogenates

Supplemental table 9: (RT)-qPCR systems

Supplemental table 10: Primer sequences used to produce overlapping amplicons for next generation sequencing

