## Supplemental data for "Favipiravir antiviral efficacy against SARS-CoV-2 in a hamster model"

### Supplemental figure 1: : *In vitro* efficacy of favipiravir

#### Summary of 50-90% drug effective concentrations and Infectious titer reductions

| | Drug effective concentration[1] (µg/mL) | | Infectious titer reduction[2] | | |
|---|---|---|---|---|---|
| | $EC_{50}$ | $EC_{90}$ | 19.6µg/mL | 39.3µg/mL | 78.5µg/mL |
| **VeroE6 cells** | | | | | |
| MOI 0.001 | 32.0 | 52.5 | 2.2 | 13.2 | 341.9 |
| MOI 0.01 | 70.0 | >78.5 | 2.0 | 5.7 | 10.9 |
| **CaCo cells** | | | | | |
| MOI 0.001 | na | na | 5.6 | 137.4 | 7720.8 |
| MOI 0.01 | na | na | 4.0 | 7.2 | 144.0 |

[1]: estimated from dose-response curves of antiviral activity (see below).
[2]: calculated using mean infectious titers without favipiravir (Virus control; see below).
MOI: multiplicity of infection. na: not applicable

#### Dose-response curves of antiviral activity in VeroE6 cells

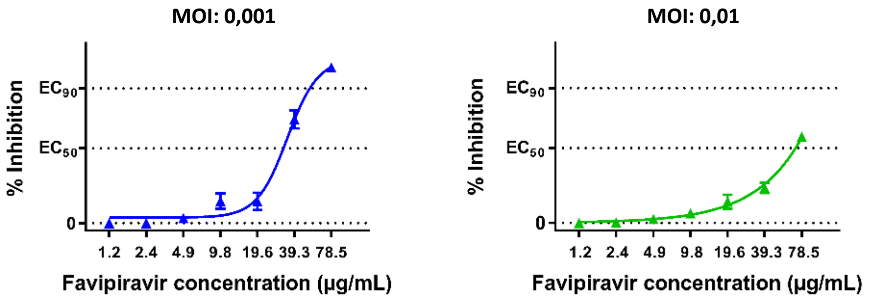


#### Infectious titer reductions

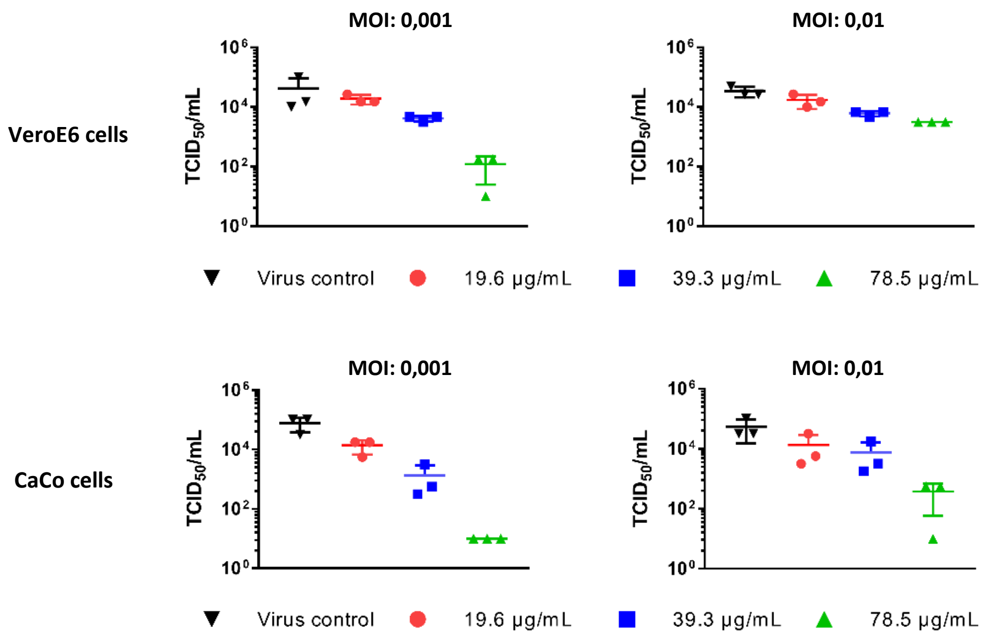

### Supplemental figure 2: Dose-response curves

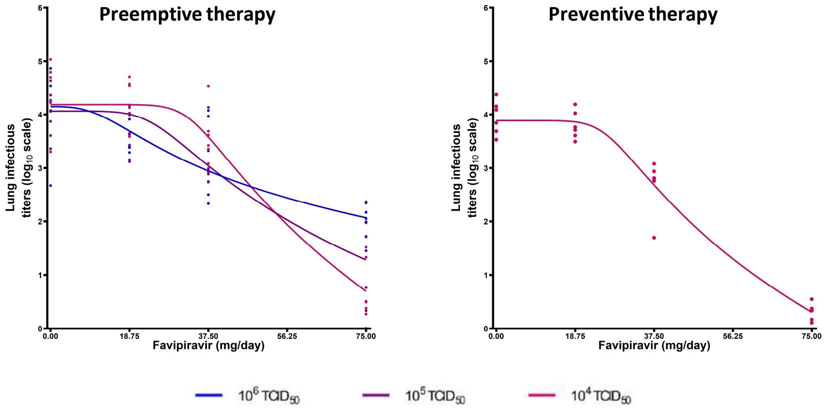

**Supplemental figure 3: Evaluation of the toxicity for animals infected and treated with high doses of favipiravir**

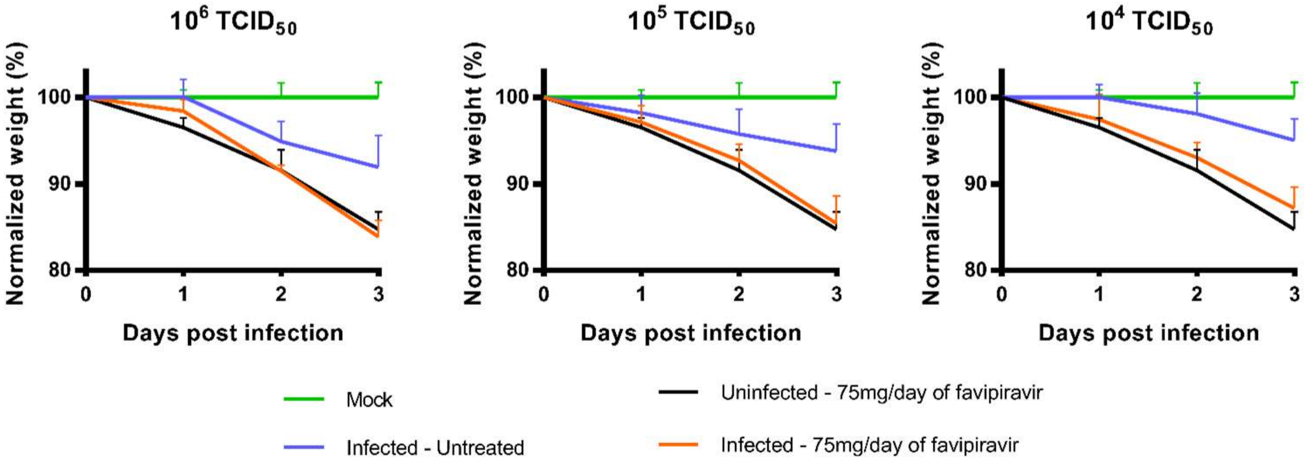


Hamsters were intranasally infected with $10^6$, $10^5$ or $10^4$ $TCID_{50}$ of virus.
Clinical follow-up with animals uninfected or infected ($10^6$, $10^5$ and $10^4$ $TCID_{50}$ of virus) and untreated or treated with a dose of Favipiravir of 75mg/day TID (preemptive antiviral therapy, see figure 2). Normalized weight at day *n* was calculated as follows: (% of initial weight of the animal at day *n*)/(mean % of initial weight for mock-infected animals at day *n*). Data represent mean ±SD. For treated animals, no significant difference was observed between uninfected and infected animals at 1, 2 and 3 dpi (Two Way ANOVA with post-hoc Sidak's multiple comparisons test).

**Supplemental figure 4: Plasma concentrations of favipiravir after administration of a single dose of favipiravir**

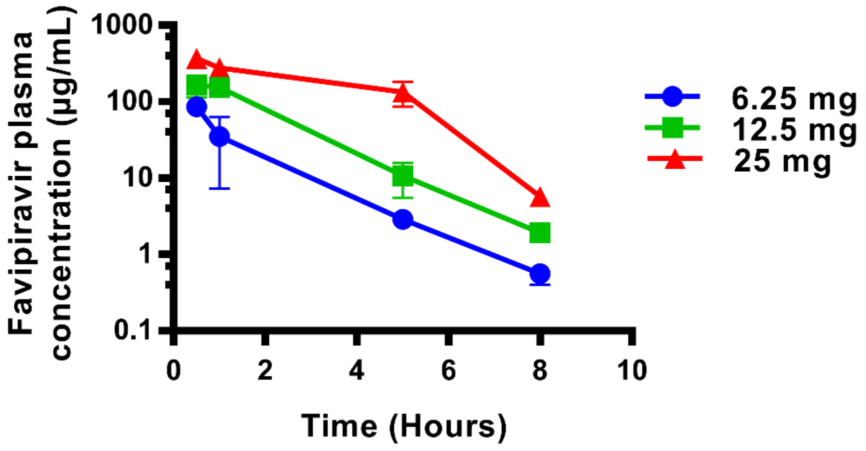


Data represent mean ±SD for three animals.

**Supplemental table 1 : Implementation of hamster model.** Normalized weights and viral RNA yields in organs.

| Inoculum ($TCID_{50}$) | Animal ID | Normalized weights | | | | | | | | Day of sacrifice | Viral RNA yields ($log_{10}$ scale) | | | Viral RNA yields (Ct values) | | | | | |
|---|---|---|---|---|---|---|---|---|---|---|---|---|---|---|---|---|---|---|---|
| | | Day 0 | Day 1 | Day 2 | Day 3 | Day 4 | Day 5 | Day 6 | Day 7 | | Lung | Plasma | Large bowel | Small bowel | Heart | Brain | Kidney | Liver | Spleen |
| | | $100 * \frac{\frac{weight\ Dn}{weight\ D_0}}{mean\ \frac{mock\ weight\ Dn}{mock\ weight\ D_0}}$ | | | | | | | | | $log_{10}\frac{Lung\ viral\ RNA\ copies}{Lung\ \gamma{-}actin\ DNA\ copies}$ | $log_{10}(Plasmatic\ viral\ RNA\ copies)$ ¤ : viral RNA yields considered at 30000cp/ml (detection threshold) | $log_{10}\frac{Large\ bowel\ viral\ RNA\ copies}{Large\ bowel\ \gamma{-}actin\ DNA\ copies}$ | | | | | | |
| $10^4$ | **253** | - | - | - | - | - | - | - | - | 2 | 4,667 | 7,357 | 0,675 | - | - | - | - | - | - |
| | **254** | - | - | - | - | - | - | - | - | 2 | 4,291 | 7,222 | 0,711 | - | - | - | - | - | - |
| | **230b** | - | - | - | - | - | - | - | - | 3 | 4,462 | 4,477 ¤ | 0,519 | - | - | - | - | - | - |
| | **229b** | - | - | - | - | - | - | - | - | 3 | 4,923 | 4,477 ¤ | 0,543 | - | - | - | - | - | - |
| | **256** | - | - | - | - | - | - | - | - | 4 | 4,010 | 4,477 ¤ | -0,220 | - | - | - | - | - | - |
| | **255** | - | - | - | - | - | - | - | - | 4 | 4,207 | 5,736 | -0,068 | - | - | - | - | - | - |
| | **201** | 100 | 98,327 | 95,601 | 90,373 | 82,362 | 76,601 | 68,920 | 68,625 | 7 | 0,739 | 4,477 ¤ | - | - | - | - | - | - | - |
| | **202** | 100 | 97,480 | 96,391 | 94,835 | 92,114 | 85,517 | 87,278 | 90,682 | 7 | 1,013 | 4,477 ¤ | - | - | - | - | - | - | - |
| | **203** | 100 | 102,636 | 102,782 | 99,562 | 92,816 | 81,871 | 81,136 | 78,877 | 7 | - | - | - | - | - | - | - | - | - |
| | **204** | 100 | 100,536 | 99,706 | 95,803 | 92,377 | 82,936 | 82,725 | 94,422 | 7 | - | - | - | - | - | - | - | - | - |
| | **205** | 100 | 98,873 | 94,540 | 93,883 | 87,948 | 77,157 | 79,843 | 85,436 | 7 | - | - | - | - | - | - | - | - | - |
| | **206** | 100 | 99,318 | 97,033 | 96,844 | 85,625 | 80,122 | 78,376 | 82,348 | 7 | - | - | - | - | - | - | - | - | - |
| | **207** | 100 | 100,920 | 99,068 | 95,340 | 93,096 | 84,128 | 76,247 | 85,088 | 7 | - | - | - | - | - | - | - | - | - |
| | **208** | 100 | 100,962 | 99,818 | 92,676 | 86,221 | 79,488 | 74,850 | 82,673 | 7 | - | - | - | - | - | - | - | - | - |
| | **209** | 100 | 100,730 | 97,358 | 95,864 | 89,786 | 78,086 | 74,320 | 79,047 | 7 | - | - | - | - | - | - | - | - | - |
| | **210** | 100 | 100,253 | 98,592 | 95,353 | 89,467 | 78,478 | 74,634 | 83,020 | 7 | - | - | - | - | - | - | - | - | - |
| | **211** | 100 | 100,116 | 97,813 | 92,199 | 88,704 | 82,012 | 85,319 | 86,916 | 7 | - | - | - | - | - | - | - | - | - |
| $10^5$ | **14** | - | - | - | - | - | - | - | - | 2 | 5,207 | 4,477 ¤ | 1,209 | - | - | - | - | - | - |
| | **13** | - | - | - | - | - | - | - | - | 2 | 5,291 | 5,799 | 2,521 | - | - | - | - | - | - |
| | **16** | - | - | - | - | - | - | - | - | 3 | 5,046 | 5,422 | -0,664 | - | - | - | - | - | - |
| | **15** | - | - | - | - | - | - | - | - | 3 | 5,075 | 5,670 | 2,295 | - | - | - | - | - | - |
| | **17** | - | - | - | - | - | - | - | - | 4 | 4,174 | 6,184 | 0,089 | - | - | - | - | - | - |
| | **18** | - | - | - | - | - | - | - | - | 4 | 4,772 | 5,546 | 0,476 | - | - | - | - | - | - |
| | **9** | - | - | - | - | - | - | - | - | 7 | 1,014 | 4,477 ¤ | -0,350 | - | - | - | - | - | - |
| | **10** | - | - | - | - | - | - | - | - | 7 | 1,554 | 4,477 ¤ | -0,194 | - | - | - | - | - | - |
| | **107** | 100 | 95,061 | 93,769 | 92,678 | 88,212 | 83,556 | 86,420 | 89,382 | 7 | - | - | - | - | - | - | - | - | - |
| | **108** | 100 | 97,611 | 90,926 | 89,244 | 82,978 | 77,279 | 80,328 | 83,817 | 7 | - | - | - | - | - | - | - | - | - |
| | **109** | 100 | 97,665 | 95,751 | 94,239 | 84,844 | 76,886 | 74,061 | 79,566 | 7 | - | - | - | - | - | - | - | - | - |
| | **110** | 100 | 95,956 | 94,543 | 91,759 | 79,057 | 74,334 | 71,738 | 76,716 | 7 | - | - | - | - | - | - | - | - | - |
| | **111** | 100 | 97,762 | 94,231 | 93,314 | 87,236 | 80,911 | 79,165 | 84,456 | 7 | - | - | - | - | - | - | - | - | - |
| | **112** | 100 | 93,307 | 91,485 | 87,593 | 79,075 | 71,175 | 69,523 | 74,126 | 7 | - | - | - | - | - | - | - | - | - |
| $10^6$ | **589** | - | - | - | - | - | - | - | - | 2 | 5,709 | 6,608 | 1,585 | no Ct | no Ct | no Ct | no Ct | no Ct | no Ct |
| | **590** | - | - | - | - | - | - | - | - | 2 | 5,653 | 5,495 | 3,092 | 34,6 | no Ct | no Ct | no Ct | 37,4 | no Ct |
| | **591** | - | - | - | - | - | - | - | - | 3 | 5,013 | 6,922 | 0,068 | no Ct | no Ct | no Ct | no Ct | no Ct | no Ct |
| | **592** | - | - | - | - | - | - | - | - | 3 | 5,271 | 6,609 | 0,310 | 36,9 | no Ct | no Ct | no Ct | no Ct | no Ct |
| | **593** | - | - | - | - | - | - | - | - | 4 | 4,412 | 6,047 | -1,731 | 31,8 | no Ct | no Ct | no Ct | no Ct | no Ct |
| | **594** | - | - | - | - | - | - | - | - | 4 | 4,523 | 6,510 | 0,567 | 32,8 | no Ct | no Ct | no Ct | no Ct | no Ct |
| | **21** | 100 | 98,616 | 95,141 | 94,398 | 89,868 | 83,065 | 82,065 | 89,252 | 7 | 0,948 | 4,477 ¤ | -0,336 | no Ct | no Ct | no Ct | no Ct | no Ct | no Ct |
| | **22** | 100 | 100,634 | 98,122 | 97,499 | 85,370 | 77,391 | 76,558 | 85,928 | 7 | 1,141 | 4,477 ¤ | -0,517 | no Ct | no Ct | no Ct | no Ct | no Ct | no Ct |
| | **19** | 100 | 102,782 | 96,754 | 97,607 | 92,486 | 84,225 | 80,989 | 89,845 | 7 | - | - | - | - | - | - | - | - | - |
| | **20** | 100 | 101,201 | 98,307 | 97,179 | 93,187 | 81,443 | 81,017 | 87,954 | 7 | - | - | - | - | - | - | - | - | - |
| | **77** | 100 | 100,881 | 96,572 | 96,067 | 85,130 | 78,221 | 81,385 | 88,226 | 7 | - | - | - | - | - | - | - | - | - |
| | **78** | 100 | 97,680 | 96,048 | 91,698 | 83,281 | 77,547 | 80,096 | 85,555 | 7 | - | - | - | - | - | - | - | - | - |
| | **79** | 100 | 98,536 | 94,483 | 93,359 | 85,585 | 75,357 | 74,562 | 82,211 | 7 | - | - | - | - | - | - | - | - | - |
| | **80** | 100 | 97,399 | 93,939 | 84,903 | 78,713 | 78,609 | 82,092 | 87,197 | 7 | - | - | - | - | - | - | - | - | - |
| Mock | **101** | 100 | 100,793 | 100,365 | 100,221 | 100,810 | 100,410 | 102,481 | 102,741 | 7 | - | - | - | - | - | - | - | - | - |
| | **102** | 100 | 101,213 | 101,999 | 102,645 | 102,637 | 102,127 | 102,876 | 102,064 | 7 | - | - | - | - | - | - | - | - | - |
| | **103** | 100 | 99,137 | 98,000 | 99,029 | 97,457 | 99,044 | 97,349 | 97,109 | 7 | - | - | - | - | - | - | - | - | - |
| | **104** | 100 | 100,148 | 100,427 | 98,540 | 97,070 | 96,823 | 96,754 | 97,172 | 7 | - | - | - | - | - | - | - | - | - |
| | **105** | 100 | 99,245 | 101,178 | 101,269 | 102,631 | 102,256 | 101,510 | 102,508 | 7 | - | - | - | - | - | - | - | - | - |
| | **106** | 100 | 99,464 | 98,030 | 98,296 | 99,395 | 99,341 | 99,031 | 98,406 | 7 | - | - | - | - | - | - | - | - | - |

**Two way ANOVA (Infected vs. Mock group)**

**Post-hoc Dunnett's multiple comparisons test**

| Day post infection | Infected hamsters | |
|---|---|---|
| | Group | Adjusted P value |
| 0 | $10^6$ $TCID_{50}$ | >0.9999 |
| | $10^5$ $TCID_{50}$ | >0.9999 |
| | $10^4$ $TCID_{50}$ | >0.9999 |
| 1 | $10^6$ $TCID_{50}$ | 0.9973 |
| | $10^5$ $TCID_{50}$ | 0.4097 |
| | $10^4$ $TCID_{50}$ | >0.9999 |
| 2 | $10^6$ $TCID_{50}$ | 0.0914 |
| | $10^5$ $TCID_{50}$ | **0.0372** |
| | $10^4$ $TCID_{50}$ | 0.5533 |
| 3 | $10^6$ $TCID_{50}$ | **0.0036** |
| | $10^5$ $TCID_{50}$ | **0.001** |
| | $10^4$ $TCID_{50}$ | **0.0131** |
| 4 | $10^6$ $TCID_{50}$ | **<0.0001** |
| | $10^5$ $TCID_{50}$ | **<0.0001** |
| | $10^4$ $TCID_{50}$ | **<0.0001** |
| 5 | $10^6$ $TCID_{50}$ | **<0.0001** |
| | $10^5$ $TCID_{50}$ | **<0.0001** |
| | $10^4$ $TCID_{50}$ | **<0.0001** |
| 6 | $10^6$ $TCID_{50}$ | **<0.0001** |
| | $10^5$ $TCID_{50}$ | **<0.0001** |
| | $10^4$ $TCID_{50}$ | **<0.0001** |
| 7 | $10^6$ $TCID_{50}$ | **<0.0001** |
| | $10^5$ $TCID_{50}$ | **<0.0001** |
| | $10^4$ $TCID_{50}$ | **<0.0001** |

### Supplemental table 3 : Statistical analysis of *in vivo* experiments.

| | Compared preemptive groups | | P value | Statistical test |
|---|---|---|---|---|
| **Lung infectious titers** | $10^6$ $TCID_{50}$ untreated | $10^6$ $TCID_{50}$ 18.75mg/day | 0.3223 | Student t-test |
| | | $10^6$ $TCID_{50}$ 37.5mg/day | 0.1099 | Student t-test |
| | | $10^6$ $TCID_{50}$ 75mg/day | **0.0001** | Student t-test |
| | $10^5$ $TCID_{50}$ untreated | $10^5$ $TCID_{50}$ 18.75mg/day | 0.2177 | Student t-test |
| | | $10^5$ $TCID_{50}$ 37.5mg/day | **0.026** | Mann whitney test |
| | | $10^5$ $TCID_{50}$ 75mg/day | **<0.0001** | Student t-test |
| | $10^4$ $TCID_{50}$ untreated | $10^4$ $TCID_{50}$ 18.75mg/day | 0.6769 | Student t-test |
| | | $10^4$ $TCID_{50}$ 37.5mg/day | **0.0384** | Student t-test |
| | | $10^4$ $TCID_{50}$ 75mg/day | **<0.0001** | Student t-test |
| **Lung viral RNA yields** | $10^6$ $TCID_{50}$ untreated | $10^6$ $TCID_{50}$ 18.75mg/day | 0.9587 | Student t-test |
| | | $10^6$ $TCID_{50}$ 37.5mg/day | 0.1038 | Student t-test |
| | | $10^6$ $TCID_{50}$ 75mg/day | **0.01** | Student t-test |
| | $10^5$ $TCID_{50}$ untreated | $10^5$ $TCID_{50}$ 18.75mg/day | 0.936 | Welch's t-test |
| | | $10^5$ $TCID_{50}$ 37.5mg/day | 0.9425 | Student t-test |
| | | $10^5$ $TCID_{50}$ 75mg/day | **0.0118** | Student t-test |
| | $10^4$ $TCID_{50}$ untreated | $10^4$ $TCID_{50}$ 18.75mg/day | 0.9372 | Mann whitney test |
| | | $10^4$ $TCID_{50}$ 37.5mg/day | 0.2231 | Student t-test |
| | | $10^4$ $TCID_{50}$ 75mg/day | **0.0001** | Student t-test |
| **Lung viral particle infectivity ratios** | $10^6$ $TCID_{50}$ untreated | $10^6$ $TCID_{50}$ 18.75mg/day | 0.0534 | Welch's t-test |
| | | $10^6$ $TCID_{50}$ 37.5mg/day | 0.1386 | Student t-test |
| | | $10^6$ $TCID_{50}$ 75mg/day | **0.0264** | Welch's t-test |
| | $10^5$ $TCID_{50}$ untreated | $10^5$ $TCID_{50}$ 18.75mg/day | 0.1333 | Student t-test |
| | | $10^5$ $TCID_{50}$ 37.5mg/day | **0.039** | Welch's t-test |
| | | $10^5$ $TCID_{50}$ 75mg/day | **0.0311** | Welch's t-test |
| | $10^4$ $TCID_{50}$ untreated | $10^4$ $TCID_{50}$ 18.75mg/day | 0.44 | Student t-test |
| | | $10^4$ $TCID_{50}$ 37.5mg/day | **0.0411** | Mann whitney test |
| | | $10^4$ $TCID_{50}$ 75mg/day | **0.019** | Mann whitney test |
| **Plasmatic viral loads** | $10^6$ $TCID_{50}$ untreated | $10^6$ $TCID_{50}$ 18.75mg/day | 0.3939 | Mann whitney test |
| | | $10^6$ $TCID_{50}$ 37.5mg/day | 0.0831 | Student t-test |
| | | $10^6$ $TCID_{50}$ 75mg/day | **0.0216** | Mann whitney test |
| | $10^5$ $TCID_{50}$ untreated | $10^5$ $TCID_{50}$ 18.75mg/day | 0.3795 | Student t-test |
| | | $10^5$ $TCID_{50}$ 37.5mg/day | 0.2879 | Mann whitney test |
| | | $10^5$ $TCID_{50}$ 75mg/day | 0.0868 | Student t-test |
| | $10^4$ $TCID_{50}$ untreated | $10^4$ $TCID_{50}$ 18.75mg/day | 0.0826 | Student t-test |
| | | $10^4$ $TCID_{50}$ 37.5mg/day | 0.1953 | Student t-test |
| | | $10^4$ $TCID_{50}$ 75mg/day | **0.0022** | Mann whitney test |

| | Compared preventive groups | | P value | Statistical test |
|---|---|---|---|---|
| **Lung infectious titers** | $10^5$ $TCID_{50}$ untreated | $10^4$ $TCID_{50}$ 18.75mg/day | 0.3989 | Student t-test |
| | | $10^4$ $TCID_{50}$ 37.5mg/day | **0.0022** | Mann whitney test |
| | | $10^4$ $TCID_{50}$ 75mg/day | **<0.0001** | Student t-test |
| **Lung viral RNA yields** | $10^5$ $TCID_{50}$ untreated | $10^4$ $TCID_{50}$ 18.75mg/day | 0.6803 | Student t-test |
| | | $10^4$ $TCID_{50}$ 37.5mg/day | **0.0233** | Student t-test |
| | | $10^4$ $TCID_{50}$ 75mg/day | **0.0022** | Mann whitney test |
| **Lung viral particle infectivities** | $10^5$ $TCID_{50}$ untreated | $10^4$ $TCID_{50}$ 18.75mg/day | 0.1767 | Student t-test |
| | | $10^4$ $TCID_{50}$ 37.5mg/day | 0.147 | Student t-test |
| | | $10^4$ $TCID_{50}$ 75mg/day | **0.0047** | One-sample t-test (hypothetical value = 0.052) |
| **Plasmatic viral loads** | $10^5$ $TCID_{50}$ untreated | $10^4$ $TCID_{50}$ 18.75mg/day | 0.613 | Student t-test |
| | | $10^4$ $TCID_{50}$ 37.5mg/day | **0.0042** | Student t-test |
| | | $10^4$ $TCID_{50}$ 75mg/day | **0.0045** | One-sample t-test (hypothetical value = 0.0049) |

### Supplemental table 4: Statistical analysis of clinical monitoring.

**Two way ANOVA (Treated vs. Mock group)**
**Post-hoc Dunnett's multiple comparisons test**

| Day post infection | Favipiravir toxicity (uninfected animals) | |
|---|---|---|
| | Group | Adjusted P value |
| 0 | 18.75mg/day | >0.9999 |
| | 37.5mg/day | >0.9999 |
| | 75mg/day | >0.9999 |
| 1 | 18.75mg/day | 0.8703 |
| | 37.5mg/day | 0.4895 |
| | 75mg/day | **0.0495** |
| 2 | 18.75mg/day | 0.9733 |
| | 37.5mg/day | 0.3462 |
| | 75mg/day | **<0.0001** |
| 3 | 18.75mg/day | 0.8776 |
| | 37.5mg/day | 0.9303 |
| | 75mg/day | **<0.0001** |
| 4 | 18.75mg/day | 0.9969 |
| | 37.5mg/day | **0.0304** |
| | 75mg/day | **<0.0001** |
| 5 | 18.75mg/day | 0.9799 |
| | 37.5mg/day | **0.0082** |
| | 75mg/day | **<0.0001** |
| 6 | 18.75mg/day | 0.9905 |
| | 37.5mg/day | 0.057 |
| | 75mg/day | **<0.0001** |
| 7 | 18.75mg/day | 0.3019 |
| | 37.5mg/day | 0.4473 |
| | 75mg/day | **<0.0001** |

**Two way ANOVA (Untreated vs. 37.5mg/day)**
**Post-hoc Sidak's multiple comparisons test**

| Day post infection | $10^5$ $TCID_{50}$ Adjusted P value | $10^4$ $TCID_{50}$ Adjusted P value |
|---|---|---|
| 0 | >0.9999 | >0.9999 |
| 1 | >0.9999 | >0.9999 |
| 2 | 0.9748 | 0,97 |
| 3 | 0.564 | 0.998 |
| 4 | 0.9989 | 0.5572 |
| 5 | 0.0126 | **<0.0001** |
| 6 | **0.0305** | **<0.0001** |
| 7 | 0.1415 | **<0.0001** |

**Supplemental table 5 : Individual data of favipiravir pharmacokinetics**

| Favipiravir pharmacokinetics in single dose (uninfected animals) | | | | | | | | | | | | | | | | |
|---|---|---|---|---|---|---|---|---|---|---|---|---|---|---|---|---|
| Favipiravir dose | Pharmacokinetics in serum | | | | | | | | Pharmacokinetics in lungs | | | | | | | |
| | T30min | | T1h | | T5h | | T8H | | T30min | | | | T5h | | | |
| | Animal ID | Favipiravir concentration (µg/ml) | Animal ID | Favipiravir concentration (µg/ml) | Animal ID | Favipiravir concentration (µg/ml) | Animal ID | Favipiravir concentration (µg/ml) | Favipiravir concentration (µg/ml) | Lung weight (mg) | Favipiravir concentration (µg/g of lung) | Lung to plasma ratio | Favipiravir concentration (µg/ml) | Lung weight (mg) | Favipiravir concentration (µg/g of lung) | Lung to plasma ratio |
| **25mg** | S | 425 | A | 336 | AB | 180 | J | 5,84 | 30,4 | 0,119 | 255 | 0,601 | 9,55 | 0,104 | 91,8 | 0,510 |
| | T | 357 | B | 258 | AC | 141 | K | 4,40 | 24,8 | 0,116 | 214 | 0,599 | 10,9 | 0,111 | 98,2 | 0,696 |
| | U | 333 | C | 243 | AD | 82,7 | L | 7,07 | 26,3 | 0,148 | 177 | 0,532 | 5,55 | 0,103 | 53,9 | 0,652 |
| **12.5mg** | V | 211 | D | 147 | AE | 15,5 | M | 2,00 | 12,48 | 0,137 | 91,1 | 0,432 | 0,74 | 0,152 | 4,87 | 0,314 |
| | W | 178 | E | 178 | AF | 11,3 | N | 1,93 | 15,48 | 0,150 | 103 | 0,581 | 0,53 | 0,117 | 4,53 | 0,401 |
| | X | 109 | F | 139 | AG | 5,25 | O | 1,89 | 8,64 | 0,111 | 77,8 | 0,715 | 0,33 | 0,155 | 2,13 | 0,406 |
| **6.25mg** | Y | 88,8 | G | 64,5 | AH | 3,10 | P | 0,74 | 7,25 | 0,132 | 54,9 | 0,618 | 0,14 | 0,13 | 1,08 | 0,347 |
| | Z | 81,5 | H | 9,10 | AI | 2,99 | Q | 0,53 | 4,28 | 0,134 | 31,9 | 0,392 | 0,13 | 0,124 | 1,05 | 0,351 |
| | AA | 88,5 | I | 31,9 | AJ | 2,62 | R | 0,42 | 8,09 | 0,127 | 63,7 | 0,720 | 0,12 | 0,105 | 1,14 | 0,436 |

| Favipiravir pharmacokinetics in multiple dose (infected animals) | | | | | | |
|---|---|---|---|---|---|---|
| Favipiravir dose | Pharmacokinetics in serum | | Pharmacokinetics in lungs | | | |
| | Ctrough (T12h J3) | | Ctrough (T12h J3) | | | |
| | Animal ID | Favipiravir concentration (µg/ml) | Favipiravir concentration (µg/ml) | Lung weight (mg) | Favipiravir concentration (µg/g of lung) | Lung to plasma ratio |
| **75mg/day** | 133 | 35,0 | | | | |
| | 134 | 29,3 | | | | |
| | 135 | 33,5 | | | | |
| | 257 | 26,5 | 1,10 | 0,106 | 10,4 | 0,39 |
| | 258 | 47,5 | 2,59 | 0,135 | 19,2 | 0,40 |
| | 259 | 36,0 | 1,93 | 0,105 | 18,4 | 0,51 |
| | 136 | 17,0 | | | | |
| | 137 | 29,1 | | | | |
| | 138 | 15,5 | | | | |
| **37.5mg/day** | 139 | 2,01 | | | | |
| | 140 | 3,06 | | | | |
| | 141 | 2,04 | | | | |
| | 260 | 4,33 | 0,18 | 0,119 | 1,51 | 0,35 |
| | 261 | 3,39 | 0,17 | 0,132 | 1,29 | 0,38 |
| | 262 | 4,02 | 0,18 | 0,142 | 1,27 | 0,32 |
| | 142 | 2,29 | | | | |
| | 143 | 0,77 | | | | |
| | 144 | 1,22 | | | | |
| **18.75mg/day** | 145 | 0,21 | | | | |
| | 146 | 0,19 | | | | |
| | 147 | 0,23 | | | | |
| | 263 | 0,40 | 0,00 | 0,132 | 0,00 | 0,00 |
| | 264 | 0,54 | 0,00 | 0,141 | 0,00 | 0,00 |
| | 265 | 0,51 | 0,00 | 0,127 | 0,00 | 0,00 |
| | 148 | 0,16 | | | | |
| | 149 | 0,35 | | | | |
| | 150 | 0,22 | | | | |

### Supplemental table 7 : Statistical analysis of mutagenic effect of favipiravir

| Analyzed data | Compared groups | Test used | p value |
|---|---|---|---|
| Total number of mutation | Favipiravir 37.5 mg/d Vs. Untreated | Unpaired t test | **0.0286 *** |
| | Favipiravir 75mg/d Vs. Untreated | Unpaired t test with Welch's correction | 0,232 |
| Total number of transition | Favipiravir 37.5 mg/d Vs. Untreated | Unpaired t test | **0.0048 **** |
| | Favipiravir 75mg/d Vs. Untreated | Unpaired t test with Welch's correction | 0,230 |
| Frequency of transition | Favipiravir 37.5 mg/d Vs. Untreated | Unpaired t test | **0.0366 *** |
| | Favipiravir 75mg/d Vs. Untreated | Unpaired t test | 0,143 |
| Total number of transversion | Favipiravir 37.5 mg/d Vs. Untreated | Unpaired t test | **0.0391 *** |
| | Favipiravir 75mg/d Vs. Untreated | Unpaired t test | 0,143 |
| Frequency of transversion | Favipiravir 37.5 mg/d Vs. Untreated | Unpaired t test | **0.0399 *** |
| | Favipiravir 75mg/d Vs. Untreated | Unpaired t test | 0,136 |
| Total number of synonymous mutation | Favipiravir 37.5 mg/d Vs. Untreated | Unpaired t test | 0,099 |
| | Favipiravir 75mg/d Vs. Untreated | Unpaired t test with Welch's correction | 0,306 |
| Frequency of synonymous mutation | Favipiravir 37.5 mg/d Vs. Untreated | Unpaired t test with Welch's correction | 0,324 |
| | Favipiravir 75mg/d Vs. Untreated | Unpaired t test | **0.0344 *** |
| Total number of non synonymous mutation | Favipiravir 37.5 mg/d Vs. Untreated | Unpaired t test | **0.0136 *** |
| | Favipiravir 75mg/d Vs. Untreated | Unpaired t test with Welch's correction | 0,200 |
| Frequency of non synonymous mutation | Favipiravir 37.5 mg/d Vs. Untreated | Unpaired t test with Welch's correction | 0,142 |
| | Favipiravir 75mg/d Vs. Untreated | Unpaired t test | **0.009 **** |
| Total number of G->A mutation | Favipiravir 37.5 mg/d Vs. Untreated | Unpaired t test with Welch's correction | **0.0068 **** |
| | Favipiravir 75mg/d Vs. Untreated | Unpaired t test with Welch's correction | 0,186 |
| Frequency of G->A mutation | Favipiravir 37.5 mg/d Vs. Untreated | Unpaired t test with Welch's correction | **0.009 ***** |
| | Favipiravir 75mg/d Vs. Untreated | Unpaired t test with Welch's correction | **0.002 **** |
| Total number of C->T mutation | Favipiravir 37.5 mg/d Vs. Untreated | Mann Whitney test | 0,057 |
| | Favipiravir 75mg/d Vs. Untreated | Mann Whitney test | 0,114 |
| Frequency of C->T mutation | Favipiravir 37.5 mg/d Vs. Untreated | Unpaired t test | 0,081 |
| | Favipiravir 75mg/d Vs. Untreated | Unpaired t test | 0,240 |

**Supplemental Table 8 : Shared mutations detected in lung clarified homogenates**

| Nucleotide position on the reference gemome NC_045512.2 | Region | Nucleotide change | Amnio-acid change | Synonymous / Non-synonymous | Transition / Transversion | Animal ID | | | | | | | | | | | | Additional information on mutation shared only within treated groups | | |
|---|---|---|---|---|---|---|---|---|---|---|---|---|---|---|---|---|---|---|---|---|
| | | | | | | Untreated | | | | Treated 37.5mg/day | | | | Treated 75 mg/day | | | | Amino-Acid mutation | Sub-localisation, if relevant | Additional information |
| | | | | | | 93 | 94 | 95 | 96 | 113 | 114 | 115 | 116 | 97 | 98 | 99 | 100 | | | |
| 373 | [ORF1ab] | G->A | E->E | Synonymous | Transition | | | | | | | | | X | | X | | - | - | - |
| 391 | [ORF1ab] | A->G | A->A | Synonymous | Transition | X | | X | X | | X | X | X | | | | | - | - | - |
| 1886 | [ORF1ab] | G->A | A->T | Non-synonymous | Transition | | | | | | X | | | | | | X | A541T | nsp2 | Original in SARS-CoV-2 (nr DB) |
| 3053 | [ORF1ab] | G->A | D->N | Non-synonymous | Transition | | | X | X | | | | | | | | X | - | - | - |
| 3743 | [ORF1ab] | C->U | H->Y | Non-synonymous | Transition | | | | | | | | X | | | X | | H1160Y | nsp3, macro domain | Original in SARS-CoV-2 (nr DB), variable residue in other CoV |
| 3990 | [ORF1ab] | C->U | T->I | Non-synonymous | Transition | | | | | X | | | | | | | X | T1242I | nsp3, SARS Unique Domain (SUD) | Original in SARS-CoV-2 (nr DB) |
| 4456 | [ORF1ab] | C->U | A->A | Synonymous | Transition | | | X | | X | | | | | | | | - | - | - |
| 5347 | [ORF1ab] | U->C | A->A | Synonymous | Transition | | | X | | | | | | | | | X | - | - | - |
| 5398 | [ORF1ab] | U->A | C->STOP | Non-synonymous | Transversion | | | X | | X | | | | | | | | - | - | - |
| 5693 | [ORF1ab] | C->U | P->S | Non-synonymous | Transition | | | | | | X | | | | | X | | P1810S | nsp3, Papain like protease (PLP) | Original in SARS-CoV-2 (nr DB), exisits also as Q at this position in SARS-CoV-2 |
| 6706 | [ORF1ab] | C->A | N->K | Non-synonymous | Transversion | | | | X | | | | | | X | | | - | - | - |
| 7083 | [ORF1ab] | C->U | S->F | Non-synonymous | Transition | | | | | | X | | | | X | | | S2273F | nsp3, Inter transmembrane domains | Substitution present in SARS-CoV-2 from Human sample (QJE38821.1) |
| 7086 | [ORF1ab] | C->U | T->I | Non-synonymous | Transition | X | | | X | | X | X | X | | | | | - | - | - |
| 8152 | [ORF1ab] | C->U | V->V | Synonymous | Transition | | | | | | X | | | | | X | | - | - | - |
| 9443 | [ORF1ab] | C->U | L->F | Non-synonymous | Transition | | | | | | | | | | | X | X | L3060F | nsp4 | Original in SARS-CoV-2 (nr DB) |
| 10070 | [ORF1ab] | A->G | M->V | Non-synonymous | Transition | X | | | | | X | | | | | | | - | - | - |
| 10738 | [ORF1ab] | U->A | N->K | Non-synonymous | Transversion | | | | | | | | | | X | | X | N3491K | nsp5 3CLpro | Original in SARS-CoV-2 (nr DB) |
| 10912 | [ORF1ab] | A->C | L->F | Non-synonymous | Transversion | X | | X | | | | | | | | | | - | - | - |
| 11273 | [ORF1ab] | G->A | V->I | Non-synonymous | Transition | | | | | X | | | | | | | X | V3670I | nsp6 | Original in SARS-CoV-2 (nr DB) |
| 11652 | [ORF1ab] | U->A | L->H | Non-synonymous | Transversion | | | | | | | | | | | X | X | L3796H | nsp6 | Original in SARS-CoV-2 (nr DB) |
| 14250 | [ORF1ab] | G->A | L->L | Synonymous | Transition | X | X | X | | X | X | X | X | X | X | | X | - | - | - |
| 18086 | [ORF1ab] | C->U | T->I | Non-synonymous | Transition | | | | X | | X | | | | | | | - | - | - |
| 18753 | [ORF1ab] | U->C | N->N | Synonymous | Transition | | | | | | X | X | | | | | | - | - | - |
| 19109 | [ORF1ab] | G->A | S->N | Non-synonymous | Transition | | | | | | | X | | | | X | | S6282N | nsp14 ExoN, N7 Mtase domain | Original in SARS-CoV-2 (nr DB) |
| 19238 | [ORF1ab] | G->A | R->K | Non-synonymous | Transition | | | | | | | | X | | | | X | R6325K | nsp14 ExoN, N7 Mtase domain | Original in SARS-CoV-2 (nr DB), R is conserved in all the analysed sequences of CoV (Ref Seq DB) |
| 19757 | [ORF1ab] | A->U | K->I | Non-synonymous | Transversion | X | | | | X | | | | | X | | | - | - | - |
| 25955 | [ORF3a] | G->A | G->D | Non-synonymous | Transition | | | | | | | | | | | X | X | G188D | - | Original in SARS-CoV-2 (nr DB) |
| 26507 | - | A->G | - | Non-coding | Transition | X | | | | | X | | | | | | | - | - | - |
| 26710 | [M] | C->U | A->V | Non-synonymous | Transition | | | | | X | X | | | | | | | A63V | - | Original in SARS-CoV-2 (nr DB) |
| 27920 | [ORF8] | C->U | I->I | Synonymous | Transition | | | | | X | X | | | | | | | - | - | - |
| 28868 | [N] | C->U | P->S | Non-synonymous | Transition | | | | | | | X | | | X | | | P199S | Linker, phosphorylation region | Original in SARS-CoV-2 (nr DB), can be T is several sequences (nr DB) |
| 29333 | [N] | A->C | N->H | Non-synonymous | Transversion | | | | X | | | | | | X | | | - | - | - |

| | |
|---|---|
| X (blue) | Shared within the group treated with 75 mg/day |
| X (purple) | Shared within the group treated with 37.5 mg/day |
| X (red) | Shared within treated groups |
| X (black) | Shared within the untreated group |
| X (green) | Present in all groups |
| X (orange) | Shared between untreated and treated with 37.5mg/day groups |
| X (cyan) | Shared between untreated and treated with 75 mg/day groups |

### Supplemental table 9: (RT)-qPCR systems

| Gene Target | Primer and probes sequences | Amplicon length | Reference |
|---|---|---|---|
| Sars-CoV-2 RNA-dependent RNA polymerase | Fwd: 5'-GTGARATGGTCATGTGTGGCGG-3'<br>Rev: 5'-CARATGTTAAASACACTATTAGCATA-3'<br>Probe: 5'-FAM-CAGGTGGAACCTCATCAGGAGATGC-TAMRA-3' | 99pb | Detection of 2019 novel coronavirus (2019-nCoV) by real-time RT-PCR (Corman et al.) |
| Syrian hamster γ-actin | Fwd: 5'-ACAGAGAGAAGATGACGCAGATAATG-3'<br>Rev: 5'-GCCTGAATGGCCACGTACA-3'<br>Probe: 5'-FAM-TTGAAACCTTCAACACCCCAGCC-TAMRA-3' | 70pb | Duplex real-time reverse transcriptase PCR to determine cytokine mRNA expression in a hamster model of New World cutaneous leishmaniasis (Espitita et al.) |
| Bacteriophage MS2 | Fwd: 5'-CTCTGAGAGCGGCTCTATTGGT-3'<br>Rev: 5'-GTTCCCTACAACGAGCCTAAATTC-3'<br>Probe: 5'-VIC-TCAGACACGCGGTCCGCTATAACGA-TAMRA-3' | 100pb | RNA and DNA Bacteriophages as Molecular Diagnosis Controls in Clinical Virology: A Comprehensive Study of More than 45,000 Routine PCR Tests (Ninove et al.) |

**Supplemental table 10: Primer sequences used to produce overlapping amplicons for next generation sequencing**

| Name | Primer Sequence | Start | End | Tm | GC% |
|---|---|---|---|---|---|
| 1Forward | ACCAACCAACTTTCGATCTCTTGT | 31 | 54 | 60.69 | 41.67 |
| 1Reverse | TTTCGAGCAACATAAGCCCGTT | 2621 | 2642 | 61.13 | 45.45 |
| 2Forward | AACAACCTACTAGTGAAGCTGTTGA | 2565 | 25989 | 60.16 | 40.00 |
| 2Reverse | TTGACATGTCCACAACTTGCGT | 5006 | 5027 | 61.26 | 45.45 |
| 3Forward | CTTCTTTCTTTGAGAGAAGTGAGGACT | 4940 | 4966 | 60.69 | 40.74 |
| 3Reverse | TGCCAAAAACCACTCTGCAACT | 7234 | 7255 | 61.47 | 45.45 |
| 4Forward | GTGGTTTAGATTCTTTAGACACCTATCCT | 7143 | 7171 | 60.59 | 37.93 |
| 4Reverse | AGGTGTGAACATAACCATCCACTG | 9644 | 9667 | 60.81 | 45.83 |
| 5Forward | ACTCATTCTTACCTGGTGTTTATTCTGT | 9558 | 9585 | 60.69 | 35.71 |
| 5Reverse | CTGGACACATTGAGCCCACAAT | 11923 | 11944 | 61.14 | 50.00 |
| 6Forward | TGCACATCAGTAGTCTTACTCTCAGT | 11864 | 11889 | 61.25 | 42.31 |
| 6Reverse | TGTGACTCTGCAGTTAAAGCCC | 14186 | 14207 | 60.81 | 50.00 |
| 7Forward | AGACGGTGACATGGTACCACAT | 13758 | 13779 | 61.41 | 50.00 |
| 7Reverse | ACACGTTGTATGTTTGCGAGCA | 15354 | 15375 | 61.63 | 45.45 |
| 8Forward | TGATTGTTACGATGGTGGCTGT | 14880 | 14901 | 60.29 | 45.45 |
| 8Reverse | GTGCAGGTAATTGAGCAGGGTC | 17437 | 17458 | 61.52 | 54.55 |
| 9Forward | TGATTTGAGTGTTGTCAATGCCAG | 17382 | 17405 | 60.26 | 41.67 |
| 9Reverse | ATTAGCAGCAATGTCCACACCC | 19845 | 19886 | 61.21 | 50.00 |
| 10Forward | AATGTAGCATTTGAGCTTTGGGC | 19774 | 19796 | 60.37 | 43.48 |
| 10Reverse | ACCAGCTGTCCAACCTGAAGAA | 22324 | 22345 | 61.82 | 50.00 |
| 11Forward | ACATCACTAGGTTTCAAACTTTACTTGC | 22263 | 22290 | 60.68 | 35.71 |
| 11Reverse | ATGAGGTGCTGACTGAGGGAAG | 24715 | 24736 | 61.74 | 54.55 |
| 12Forward | GTCAGAGTGTGTACTTGGACAATCA | 24649 | 24673 | 60.74 | 44.00 |
| 12Reverse | ACTGCTACTGGAATGGTCTGTGT | 27142 | 27164 | 61.58 | 47.83 |
| 13Forward | GGTGACTCAGGTTTTGCTGCAT | 27087 | 27108 | 61.65 | 50.00 |
| 13Reverse | CGTAAACGGAAAAGCGAAAACGT | 29571 | 59593 | 61.08 | 43.48 |

The primers used come from a larger series of primer intended initially for carrying out multiplexed PCR obtained on the site http://primal.zibraproject.org/, acording to Quick J et al. Multiplex PCR method for MinION and Illumina sequencing of Zika and other virus genomes directly from clinical samples. Nat Protoc. 2017 Jun;12(6):1261-1276.
